# Chromatin remodelling enables enhancer resetting to facilitate the ERK transcriptional response

**DOI:** 10.64898/2026.03.27.714694

**Authors:** Ramy Ragheb, Nicola Reynolds, Devina Shah, Maya Lopez, Jennifer Balmer, Nikolaos Markozanis, Prajakta Gade, Andria Koulle, Oluwaseun Ogundele, Robin Floyd, Ernest Laue, Brian Hendrich

**Affiliations:** Living Systems Institute, University of Exeter, Exeter EX4 4QD, UK; Cambridge Stem Cell Institute, Jeffrey Cheah Biomedical Centre, University of Cambridge, Cambridge CB2 0AW, UK; Department of Biochemistry, University of Cambridge, Cambridge CB2 1GA, UK; University of Copenhagen, Novo Nordisk Foundation Center for Protein Research, Blegdamsvej 3B, 2200 Copenhagen, Denmark; Department of Life Sciences, Imperial College London, London, SW7 2AZ, UK; Centre for Biodiversity Genomics, University of Guelph, 50 Stone Rd E, Guelph, ON, N1G 2W1, Canada

## Abstract

During development, cellular identity is ultimately determined by transcriptional output: lineage-specific genes must be activated, while genes associated with alternative fates must be repressed. This process depends on the activity of chromatin remodelling complexes, which regulate the accessibility of transcription factors to chromatin regulatory elements. In addition, cellular identity is shaped by exposure to intercellular signals. Understanding the mechanisms by which extracellular signals are translated into changes in the transcriptional program is essential for understanding cell fate decisions during development, as well as in disease conditions such as cancer. Here we describe a rapid and widespread enhancer resetting event in response to ERK signalling in mouse ES cells. This process occurs in two distinct phases: an immediate, genome-wide alteration in transcription factor binding dynamics at regulatory regions which is dependent on the release of paused RNA Polymerase II, followed by the re-establishment of a context appropriate, stable chromatin state. We demonstrate that the chromatin remodelling complex NuRD is required for this reestablishment phase and for the appropriate transcriptional response to ERK signalling. We propose that enhancer resetting places genomic regulatory regions in a state which is permissive to the exchange of transcription factors in order to establish a new, stable enhancer topology enabling rapid yet precise transcriptional response to extracellular signals.

## Introduction

During the course of development cells must respond to extracellular cues imparted by their environment. Appropriate cellular responses are vital to ensure development proceeds normally, whereas inappropriate responses, or a failure to respond, can result in developmental defects and/or oncogenesis. A cell responding to instructional cues must rapidly change its gene expression programme. This requires substantial remodelling of chromatin and of the topological organisation of key regulatory sequences: enhancers and promoters.

In mouse ES cells the ERK pathway drives exit from naïve pluripotency, the first step towards lineage commitment^1,2^. Extracellular stimulation in ES cells is driven by autocrine production of Fgf4, which switches on a phospho-relay pathway resulting in rapid phosphorylation and activation of the ERK1 and ERK2 kinases^3^. These kinases then phosphorylate a large number of substrates, including other kinases, which both amplifies the signal, resulting in a peak of ERK activity after about 40 minutes, and also induces autoregulatory feedback loops, resulting in cyclic waves of ERK signalling with a period on the order of several hours^4^. Phosphorylation targets of ERK pathway kinases include transcription factors (TFs), chromatin proteins and RNA Polymerase II^5–8^, indicating that a key function of the ERK signalling pathway is to directly impact on chromatin and transcription.

How ERK pathway stimulation drives immediate early gene expression has been extensively studied. ERK pathway kinases phosphorylate histone H3 on residues S10 and S29. This stimulates acetylation of adjacent lysine residues (K9 and K27) which then leads to further histone acetylation and recruitment of Mediator and RNA Polymerase II to promoter sequences which facilitate transcription^9–13^. ERK activation also stimulates phosphorylation of histone variant H3.3 on residue S31 in gene bodies, stimulating H3K36 methylation and facilitating transcriptional elongation^14^. Exactly how ERK signalling stimulates enhancer activity, which ultimately determines which genes respond to signals, is not clear.

Chromatin remodelling proteins play a pivotal role in deciding whether and how pluripotent embryonic stem (ES) cells respond to developmental cues. These complexes regulate gene expression by modulating the accessibility of key regulatory regions. For example, the BRG1/BRM-associated factor (BAF) complex promotes chromatin accessibility by decreasing nucleosome occupancy, whereas the Nucleosome Remodelling and Deacetylase complex (NuRD) complex restricts access at these loci. NuRD is comprised of a chromatin remodelling subcomplex made up of the ATPase/nucleosome remodeller subunit CHD4 along with GATAD2A/B and DOC1 (CDK2AP1), and a deacetylation subcomplex containing the lysine deacetylases HDAC1 and/or HDAC2 as well as a combination of the MTA1/2/3 and RBBP4/7 proteins. These two subunits are held together by the MBD3 protein^15,16^. NuRD’s chromatin remodelling activity is required for ES cells to appropriately change their gene expression programmes and undergo lineage commitment in response to differentiation cues^17–21^. In contrast the histone deacetylase activity reflects transcriptional status and is likely to be important for reinforcing transcriptional states^21,22^.

Here we sought to precisely determine how the MEK/ERK extracellular signalling pathway and NuRD’s remodelling activity work together to induce transcriptional change. We found that ERK activation results in a dramatic change in transcription factor binding kinetics at enhancer chromatin within 10 minutes, with binding kinetics returning to pre-signal levels from 30 minutes to 4 hours post signal. This change is dependent on release of paused RNAPII and coincides with a change in nucleosome positioning within enhancers and with transient changes in global enhancer chromatin accessibility. We refer to this fast, but transient change in TF-enhancer interactions as “Enhancer Resetting.” Using acute depletion of two different NuRD components, and depletion or inhibition of the catalytic component of the BAF complex, we show that while the initial change in TF binding kinetics is NuRD and BAF independent, the subsequent recovery of TF binding profiles, as well as the immediate changes in gene expression, is dependent on chromatin remodeller activity. We therefore define the immediate chromatin response to activation of a major signalling pathway, which involves a previously unidentified and chromatin remodeller-dependent Enhancer Resetting event.

## Results

### ERK activation causes a transient change in transcription factor binding kinetics

Mouse ES cells are maintained in a naïve state using defined media which includes small molecule inhibitors of MEK/ERK signalling (PD0325901) and GSK3 (CHIRON/CHIR99021) and the cytokine LIF (Leukaemia Inhibitory Factor), called “2iLIF” media (Figure 1A)^23,24^. This defined media allows for precise modulation of these signalling pathways simply by removing individual inhibitors. Crucially, mouse ES cells can be maintained in culture short-term in the presence of just the GSK3 inhibitor with LIF (1iLIF, Figure 1A) thus activating the ERK pathway without resulting in a change of cell state^23^. Autocrine ERK pathway activation is first detectable as the appearance of phosphorylated ERK between 5-10 minutes of withdrawal of the PD0325901 inhibitor (PD03) (Figure 1B).

**Figure 1.**
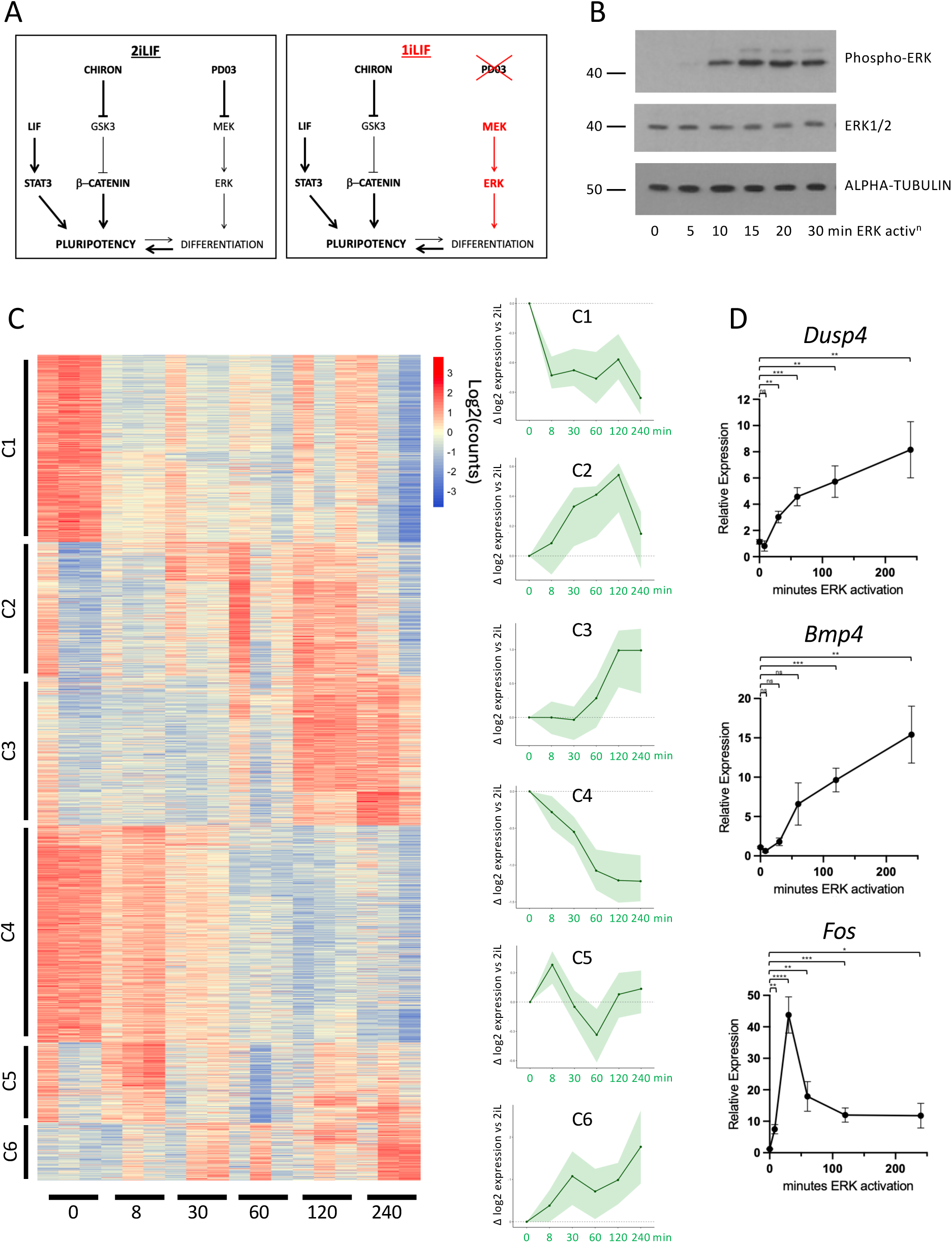
ERK Activation rapidly changes gene expression in mouse ES cells. **A**. Schematic of the two inhibitors with LIF (2iLif) culture condition for maintenance of naïve ES cells. The cytokine LIF and the GSK3 inhibitor CHIR99021 (Chiron) both promote pluripotency, while the MEK/ERK inhibitor PD0325901 (PD03) blocks differentiation (left panel). If the MEK/ERK inhibitor is removed (right panel), the MEK/ERK pathway is active which promotes expression of differentiation-associated genes. However the combined action of LIF and CHIRON keep ES cells in an undifferentiated state. **B**. Western blot of ES whole cell extracts across the ERK activation time course, with alpha-tubulin and total ERK1/2 acting as loading controls (bottom panels). Pathway activation is indicated by the presence of phosphorylated ERK (top panel). The location of size markers in kDa is indicated at left. **C**. Heatmap of nuclear RNA-seq carried out before (in 2iL conditions, 0 timepoint) and after indicated times of ERK activation (bottom, minutes). Three biological replicates for each time point are shown, and the cumulative behaviour of each gene cluster (indicated at left) is shown in a cumulative plot on the right. Line shows mean gene expression; shaded ribbon indicates the interquartile range (25th–75th percentile). **D**. qPCR data for indicated genes across the ERK activation time course. Mean expression of N = 4 biological replicates with standard error is plotted. * p<0.05, ** p< 0.01, *** p < 0.001, **** p < 0.0001 by two-tailed t-test.

The ultimate consequence of ERK pathway activation in ES cells is a change in gene expression. RNAseq using nuclear RNA to enrich for nascent transcripts across a time course of ERK activation showed that while changes in transcription were evident from as early as 8 minutes after inhibitor withdrawal (Figure 1C, Clusters 1, 4, 5) most ERK target genes were not induced until later, between 30 and 60 minutes (Figure 1C, Clusters 2, 3, 6). The kinetics of these transcriptional changes were confirmed using both RT-qPCR for nascent transcripts (Figure 1D) and nascent RNA sequencing (4SU-RNAseq; Figure S1). Transcription of immediate early genes such as *Fos* occurred rapidly, within 8 minutes of inhibitor removal (Cluster 2 in Figure 1C and Figure 1D). Transcription of known ERK target genes such as *Dusp4* and *Bmp4* were significantly increased later, from 30 and 60 minutes respectively, and continued to increase through 4 hours (Cluster 3 in Figure 1C and Figure 1D). This indicates that transcriptional activation starts during the first wave of ERK activity and is then sustained through subsequent cycling of ERK activation. Furthermore, while signal activation is very rapid (≤10 minutes), the resulting changes to chromatin and/or the transcriptional machinery at most ERK responsive genes require an additional 20 - 30 minutes to impact transcriptional output. This window, between signal activation and transcriptional response, allowed us to identify chromatin changes that direct transcription in response to extracellular signals.

Enhancers are thought to alter the activation kinetics of RNA polymerase II^25^ so we first asked whether ERK activation induced an immediate change in the levels of RNA Polymerase II at responsive enhancers. Chromatin immunoprecipitation (ChIP) for either total RNA Polymerase II (POLR2A) or for the Serine 5 phosphorylated form in formaldehyde (FA)-fixed cells showed no detectable change in enrichment at an ERK-responsive enhancer located ∼23 Kb upstream of the *Bmp4* gene after 8 or 30 minutes of ERK activation (Figure 2A). Levels of both forms were increased by 4 hours, consistent with increased transcription of *Bmp4*. Similarly, levels of both the Mediator complex subunit, MED1, and acetylated H3K27 (H3K27Ac) increased coincident with increased transcription rather than before (Figure 2A, Figure S2A). As we found that major changes in the levels of total or active RNA Polymerase II, MED1 and acetylation of H3K27 occurred after the 30 minute window that precedes transcriptional activation, it is unlikely that pre-emptive accumulation of these factors drive ERK-dependent transcriptional change.

**Figure 2.**
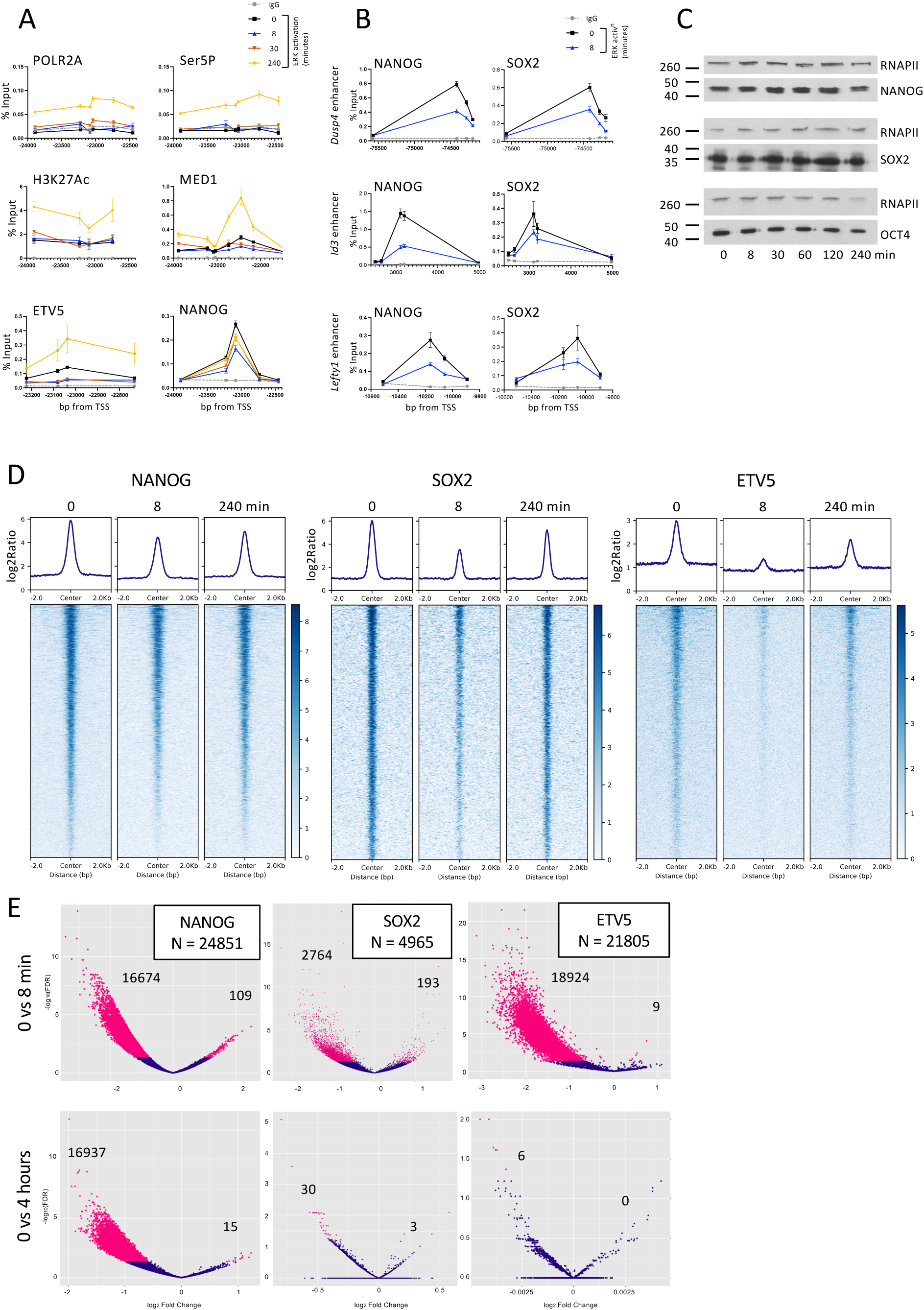
ERK activation results in a rapid but transient change in transcription factor ChIP-seq enrichment across enhancers. **A**. FA-fixed ChIP-qPCR data for indicated proteins at indicated times of ERK activation across an enhancer located ∼23 Kb upstream of the *Bmp4* locus. X-axis labels indicate base pairs relative to the annotated *Bmp4* TSS. Mean ± standard error is plotted from ≥3 biological replicates. **B**. FA fixed ChIP-qPCR data for NANOG or SOX2 across the indicated enhancers in 2iL conditions (black) or after 8 minutes of ERK activation (blue). X-axis shows base pairs relative to the annotated TSS of each gene. Mean ± standard error is plotted from ≥3 biological replicates. **C**. Western blots of nuclear extracts collected at indicated times of ERK activation (bottom) and probed with antibodies for proteins indicated at right. The position of size markers in kDa is indicated at left. **D**. Heatmaps of FA fixed ChIP-seq carried out for NANOG, SOX2 or ETV5 either before (0) or at indicated times after of ERK activation. N = 3 biological replicates each. **E**. Volcano plots of all peaks identified for NANOG, SOX2 or ETV5 showing relative change in enrichment between 0 and 8 minutes (top panels) or 0 and 240 minutes (bottom panels). Numbers of peaks showing significant changes (red points) are indicated in the figure.

We next asked whether a change in transcription factor binding could drive transcriptional change. ETV5 is an ERK-responsive transcription factor which is important for ES cells to exit the naïve state^26^. We therefore tested whether early accumulation of ETV5 at enhancers might drive the ERK-dependent transcriptional response.

Similar to our findings for transcription machinery and H3K27 acetylation, we observed no increase in ETV5 levels at the *Bmp4* enhancer within 30 minutes of ERK activation (Figure 2A). Enrichment of ETV5 did however increase by 4 hours of activation, coincident with increased expression at this locus. Contrary to our expectations, ETV5 enrichment decreased relative to starting levels after 8 and 30 minutes of ERK activation, an effect which was also seen for MED1 (Figure 2A). To investigate this apparent loss of binding further, we followed changes in the enrichment of two pluripotency-associated transcription factors, NANOG and SOX2, at enhancers associated with ERK-responsive genes. NANOG, like ETV5 and MED1, showed an immediate but transient decline in enrichment at the *Bmp4* enhancer after 8 minutes of ERK activation (Figures 2A, S2B). This acute loss of enrichment was also seen for both NANOG and SOX2 at enhancers located near the ERK-responsive *Dusp4*, *Id3* and *Lefty1* genes (Figures 2B, S2B). (No enrichment for SOX2 at the *Bmp4* enhancer was detected in any conditions.) We detected no evidence for a change in the steady-state levels of either of the transcription factors across this time course, nor evidence for ERK-induced phosphorylation of NANOG or SOX2 as judged by apparent sizes on western blots (Figure 2C), in agreement with phosphoproteomic and immunofluorescence studies^5,6^.

To assess the extent of this change in transcription factor behaviour, we performed calibrated ChIP-seq for NANOG, SOX2 and ETV5 in FA-fixed cells across the ERK activation time course. Consistent with our ChIP-qPCR results, enrichment of all three transcription factors was reduced across enhancers globally after 8 minutes of ERK activation relative to 2iLIF levels, showing signs of recovery by 4 hours (Figure 2D). The magnitude of this effect varied between transcription factors. Following ERK activation, just over half of SOX2 peaks identified in 2iLIF conditions showed reduced enrichment (2764 of 4965; 55%), a larger proportion of NANOG peaks were affected (16,674 of 24,851; 67%) while the strongest effect was observed for ETV5, with over 85% of peaks (18,924 of 21,805) showing significantly reduced enrichment after 8 minutes (Figure 2E).

For both ETV5 and SOX2, very few peaks exhibited significant differences in enrichment between 2iLIF and 4 hour conditions, indicating near-complete restoration of their enrichment profiles (Figure 2D, E). Although NANOG also showed some recovery of enrichment after 4 hours (Figure 2D) levels remained lower than those observed under 2iLIF conditions and a large number (16,937 of 24,851; 68%) of peaks were differentially enriched (Figure 2E; note X-axis scales). Overlap between regions showing reduction in ChIP signal at 8 minutes for the three transcription factors was relatively minor suggesting either transcription factor – locus specificity or a more stochastic effect (Figure S2C). Although we can detect no change in NANOG protein levels by western blotting across this time course (Figure 2C), NANOG levels have been shown to begin to decrease from approximately 6 hours post-ERK activation using more quantitative methods^27^. It is therefore possible that the reduced NANOG enrichment we see at 4 hours is due to a small but significant decrease in levels of nuclear NANOG protein.

Enhancers showing differential association with NANOG, SOX2 and ETV5 at 8 minutes displayed hallmarks of active enhancers (high H3K4Me1 and H3K27Ac, low H3K4Me3), but very little binding was detected at inactive enhancers (high H3K4Me1, lowH3K27Ac and H3K4Me3) (Figure S2D). This transient, ERK-induced change in transcription factor behaviour therefore does not convert inactive enhancers to active enhancers; rather it occurs predominantly or exclusively on enhancers bearing histone marks associated with active chromatin.

Crosslinking of proteins and chromatin by formaldehyde is inefficient for highly dynamic interactions, however these interactions can be detected if cells are crosslinked with DSG (disuccinimidyl glutarate) prior to FA fixation^28–30^. To distinguish whether the observed loss of ChIP signal was caused by a loss of transcription factor occupancy or by a change in binding kinetics, we repeated the ChIP on cells double-fixed with both DSG and FA. In stark contrast to the results obtained with FA-fixed cells, ChIP-qPCR on double-fixed chromatin showed either a gain or no significant change in transcription factor enrichment for NANOG, ETV5 and SOX2 at the 8-minute time point (Figure 3A). These fixation-dependent ChIP results are therefore likely to be caused by some change in transcription factor behaviour, and do not reflect abundance on chromatin.

**Figure 3.**
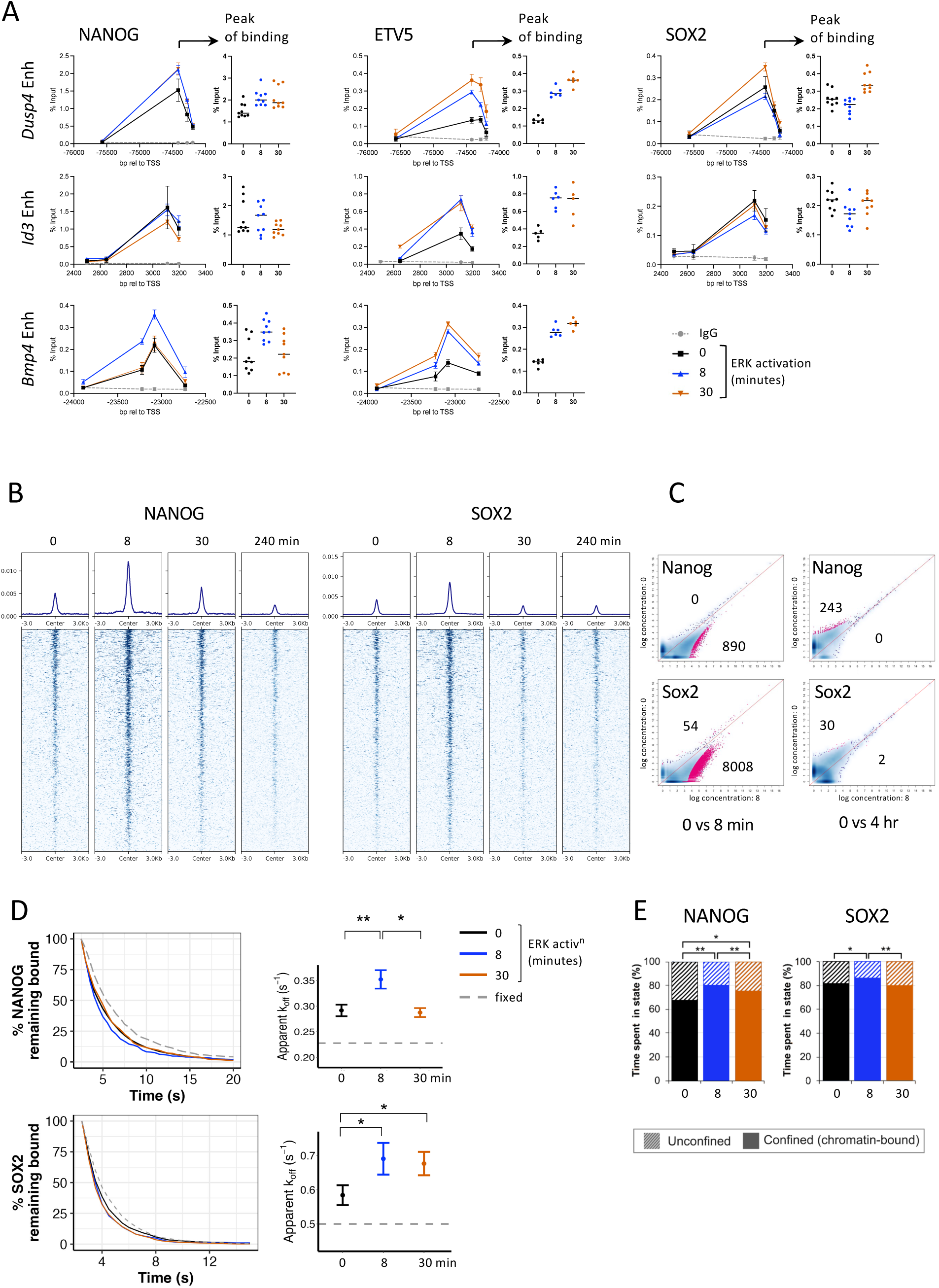
ERK activation induces a change in transcription factor binding kinetics. **A.** ChIP-qPCR data in DSG/FA fixed cells is plotted for indicated transcription factors (top) at indicated loci (left) either before (0) or after 8 or 30 minutes of ERK activation. In each case all replicates for the peak of binding are re-plotted at right for indicated time points. Line drawings show mean and standard error (N ≥ 2) while smaller plots show all replicate values at peak of binding and the horizontal line indicates the mean. **B**. Heatmaps of Cut&Run data for NANOG or SOX2 at indicated time points of ERK activation. N = 3 biological replicates. **C**. Enrichment of peaks identified at time 0 are compared to enrichment at 8 minutes (left) or 4 hours (right). Significantly changing peaks are coloured in red, and the number of peaks significantly losing enrichment in each comparison is indicated below the diagonal, while the number gaining enrichment is indicated above the diagonal. **D**. Fluorescence survival curves of chromatin-bound NANOG-HALO (upper left) and SOX2-HALO (lower left) localisations in 2iL and upon ERK activation. Apparent dissociation rates (k_off_) of chromatin-bound NANOG-HALO (upper right) and SOX2-HALO molecules (lower right) were calculated through fitting a single exponential decay model to the survival curves at left. Mean and 95% confidence intervals for each fit, applied to data taken from four independent experiments, are plotted at right. The horizontal dashed line represents the upper 95% confidence limit for a fixed-cell control. * indicates that 95% confidence intervals do not overlap, while ** indicates that 99% confidence intervals do not overlap. **E**. Confined fractions of NANOG-HALO (left) and SOX2-HALO (right) in 2iL and upon ERK activation in ChL media as determined from 3D short-exposure Single Molecule Tracking. Significance was assessed by Chi-square proportion test on data from three independent experiments. *p<0.5, **p<0.01.

To obtain a more accurate global picture of TF binding during ERK activation, we turned to Cut&Run, which does not require the use of crosslinking and hence is not subject to artefacts introduced by chromatin fixation^31^. Cut&Run for NANOG and SOX2 showed an increase in signal after 8 minutes of ERK-activation at binding sites genome-wide (Figure 3B, C), in contrast to our FA-fixed ChIPseq data. This further supports our assertion that TF binding kinetics underlie the observed change in FA-ChIP enrichment. While SOX2 binding appears to change little between 0 minutes and 4 hours, NANOG Cut&Run enrichment continued to decline through 4 hours, consistent with our ChIP-seq results (Figures 3B, C compared to Figure 2). This likely indicates that the loss of enrichment seen in NANOG ChIP-seq at the latest time points reflects a genuine decrease in NANOG binding at enhancers 4 hours after ERK activation, possibly due at least in part to reduced NANOG protein levels. We therefore propose that the rapid loss of transcription factor enrichment seen in FA-fixed cells, but not in double-fixed or unfixed cells, is most likely due to an ERK-induced change in the binding kinetics of transcription factors on enhancer chromatin globally. Specifically, we predict that ERK activation leads to an acute decrease in the residence time, and hence an increase in the dissociation rate of transcription factors at enhancer chromatin.

To test this prediction, we directly measured transcription factor binding kinetics using single molecule imaging in live cells. Endogenous *Nanog* or *Sox2* genes were targeted to create heterozygous C-terminal HALO-tag fusions. Using a double helix point-spread function microscope^32^ we generated 3D tracks of individual HALO-tagged NANOG or SOX2 proteins in the nucleus. Using a 500 ms exposure resulted in motion blurring of freely diffusing proteins, allowing us to focus on the slower moving chromatin-bound proteins^33^. In 2iLIF conditions the apparent dissociation rate (K_off_) for NANOG was calculated as 0.29 ± 0.011 s^-1^, while after 8 minutes of ERK activation the dissociation rate increased to 0.35 ± 0.018 s^-1^. Similarly, K_off_ for SOX2 in 2iLIF conditions was 0.58 ± 0.029 s^-1^, which increased to 0.69 ± 0.046 s^-1^ after 8 minutes of ERK activation (Figure 3D). While the NANOG K_off_ returned to near normal levels by 30 minutes (0.28 ± 0.009 s^-1^), that of SOX2 did not decrease significantly by 30 minutes (Figure 3D). The change in K_off_ at 8 minutes was accompanied by an increase in the fraction of stably bound protein for both NANOG and SOX2 (Figure 3E), consistent with the increase in enrichment seen for both proteins by Cut&Run at 8 minutes (Figure 3C). Together these data show that ERK signal activation leads to an acute change in transcription factor binding kinetics. For both NANOG and SOX2 the proportion of bound protein increases, but their dissociation rates also increase, meaning that each individual protein spends less time bound to DNA and hence is less efficiently cross-linked to chromatin upon formaldehyde treatment. These data are consistent with a model in which ERK activation causes an immediate but transient destabilisation of transcription factor binding to regulatory elements followed by restoration, which we call “Enhancer Resetting”.

### Pause release of RNAPII destabilises TF-chromatin interactions to initiate Enhancer Resetting

Enhancer Resetting consists of a rapid mobilisation of transcription factor-chromatin interactions across all active enhancers in response to a signalling event. We hypothesised that such an immediate and widespread effect would likely involve a factor which occupies these loci in the steady state. The transcription machinery is present at low levels at all active enhancers and would be a key target for regulation. We therefore asked whether Enhancer Resetting, specifically the initial TF mobilisation step, was dependent on RNA Polymerase II activity. To this end we employed specific small molecule inhibitors to block different aspects of RNA Polymerase II activity: BAY299 to inhibit pre-initiation complex (PIC) formation; Triptolide to prevent transcription initiation and DRB to prevent release of paused RNA Polymerase^34,35^. Cells were cultured in the presence of inhibitor for 1 hour prior to ERK activation and formaldehyde fixation. Inhibition of either PIC formation or transcription initiation had no impact on Enhancer Resetting, evident as a reduction in NANOG ChIP signal after 8 minutes of ERK activation in formaldehyde fixed samples (Figure 4A). This indicates that the change in TF binding kinetics does not require either recruitment of new RNA polymerase or new transcription initiation. In contrast, inhibition of pause release resulted in a reduction in NANOG signal in steady state 2iLIF conditions, but no further decrease upon ERK activation (Figure 4A). This shows that release of paused RNAPII – i.e. significant transcription through enhancers – is important for the change in TF binding kinetics characteristic of Enhancer Resetting.

**Figure 4.**
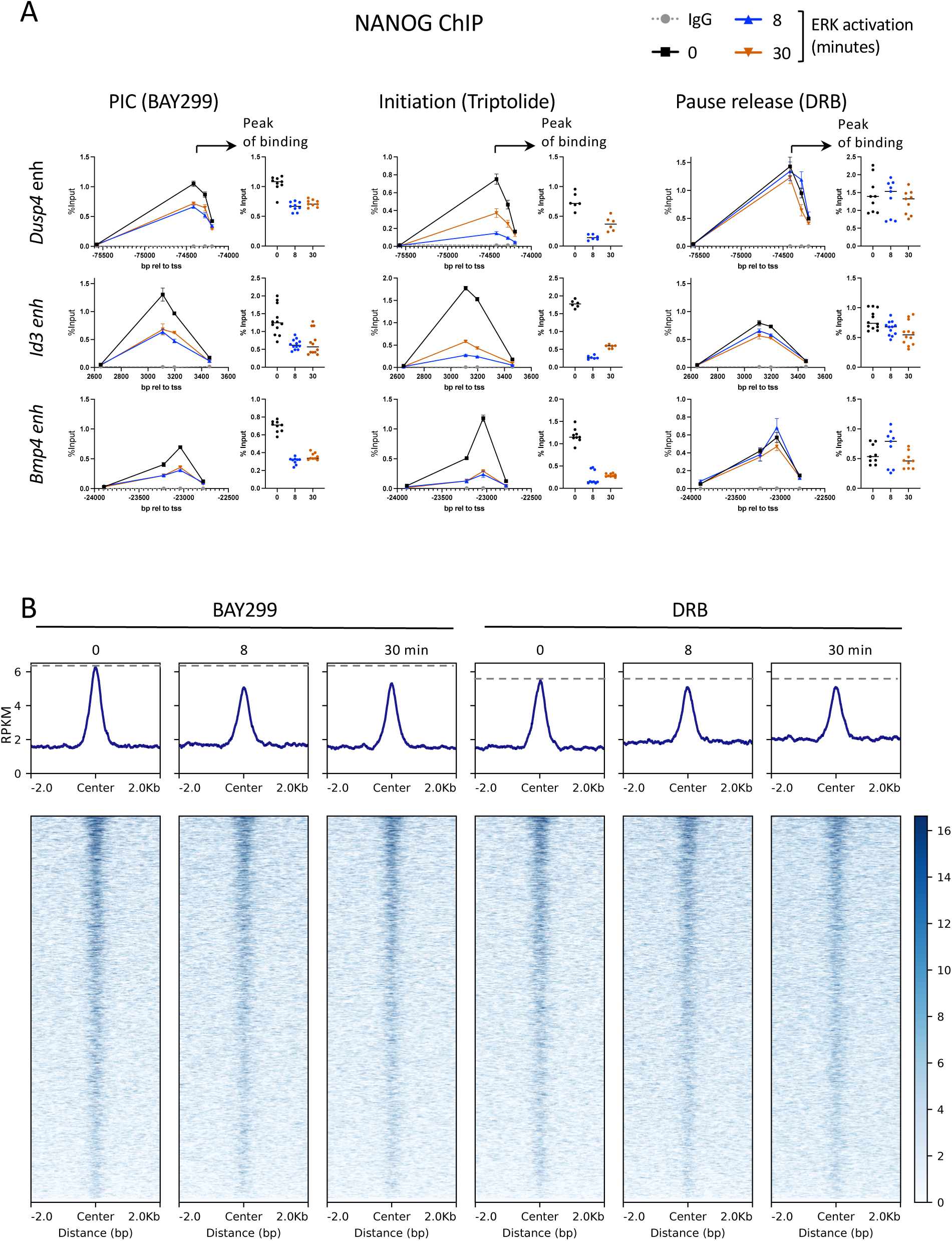
Release of RNA polymerase pausing is required for the initiation of Enhancer Resetting. **A.** FA fixed ChIP-qPCR is plotted as in 3A in the presence of indicated RNA Polymerase II inhibitors. N ≥ 2 biological replicates. **B**. Heatmaps of FA fixed ChIP-seq for NANOG carried out in the presence of indicated inhibitors either before (0) or after 8 or 30 minutes of ERK activation. Level of enrichment in starting condition is marked with dashed line for ease of comparison between timepoints. N = 3 biological replicates.

We confirmed this genome-wide by performing ChIPseq for NANOG in cells treated with either BAY299 or DRB before ERK activation. NANOG signal across active enhancers was reduced after 8 minutes of ERK activation in the presence of BAY299, confirming that new transcription initiation is not required for enhancer resetting (Figure 4B). Blocking pause release with DRB, however, prevented this decrease in NANOG enrichment across its binding sites, confirming that release of paused RNAPII at active enhancers globally drives the rapid and widespread destabilisation of TF binding in response to ERK signalling in the initial step in enhancer resetting (Figure 4B).

### ERK activation causes a transient change in enhancer chromatin

Transcription factor binding kinetics are known to be influenced by nucleosome density^36,37^, so we next used ATAC-seq to ask whether the dramatic changes in TF behaviour seen upon ERK activation coincided with changes in chromatin structure. We focussed on those loci which are clearly subject to Enhancer Resetting, i.e. those bound by TFs in 2iLIF conditions where these interactions are destabilised after 8 minutes of ERK activation. These are regulatory regions that we expect to be active in 2iL conditions and therefore their chromatin should be accessible at the start of the time course. Nevertheless, there was an immediate increase in the number of Tn5 integrations at resetting regions for all three TFs, indicating a significant increase in accessibility induced by ERK activation (Figure 5A). This ERK-induced increase in accessibility was transient as integration frequencies decreased to below starting (2iLIF) levels by 30 minutes and 4 hours, reflecting a subsequent reduction in accessibility. ERK activation thus results in a transient increase in chromatin accessibility at TF-bound sequences which correlates with reduced TF residence times.

**Figure 5.**
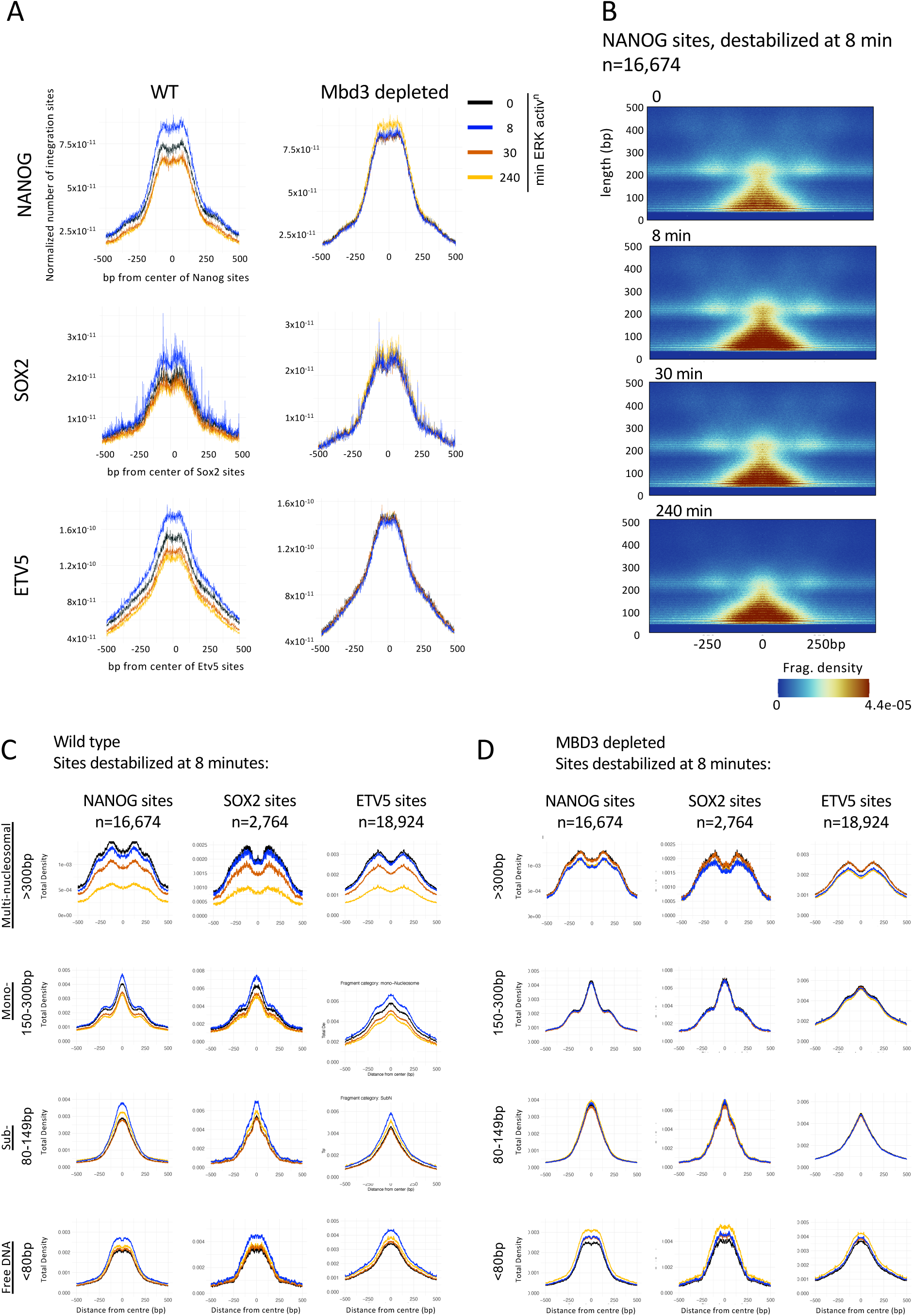
Dynamic chromatin changes occurring during Enhancer Resetting require an intact NuRD complex. **A**. Tn5 integration frequency was determined from ATAC-seq after indicated times of ERK activation and plotted across sites showing a change in FA-ChIPseq enrichment for indicated transcription factors after 8 minutes of ERK activation. Profiles are shown in wild-type cells (left panels) or after MBD3 depletion (right panels). **B**. Volcano plots constructed from ATAC-seq data across sites showing changes in NANOG binding behaviour after 8 minutes of ERK activation in wild type cells. **C**. Density of fragments of indicated sizes obtained from Vplot data (see Methods) plotted across sites showing acute changes in binding for NANOG, SOX2 or ETV5. **D**. As in **C** but for cells depleted for MBD3.

To assess whether this increase in accessibility corresponded to a change in nucleosome organisation, we constructed Vplots of the ATAC-seq data^38,39^. These showed a decrease in larger fragments (>300 bp) by 4 hours for NANOG, ETV5 and SOX2 target sites, as well as dynamic changes in the abundance of sub-nucleosome sized (<200 bp) reads (Figure 5B, Figure S3). To better quantify chromatin changes we plotted the density of fragment lengths representing multi-nucleosomes (>300 bp), single nucleosomes (150 – 300 bp), partial nucleosomes (80 - 149 bp) and free DNA (<80 bp). This analysis showed that ERK activation results in an immediate loss of multi-nucleosome sized reads, with a corresponding gain in mono-nucleosome, sub-nucleosome and nucleosome-free sized reads at responsive enhancers (Figure 5C). While the decrease in multi-nucleosome reads continues progressively across the time course, the increase in mono-nucleosome sized reads is transient and is followed by a loss of these reads at later time points to levels below that seen in 2iLIF. Reads corresponding to subnucleosomes and nucleosome free regions also decrease after 8 minutes, but remain close to, or increased relative to levels seen in 2iLIF conditions. We postulate that ERK activation induces a rapid decrease in nucleosome density within these open chromatin regions, such that the probability of generating multi-nucleosome reads from Tn5 integrations is immediately reduced, while that of generating single and sub-nucleosome sized reads increases. This event happens very quickly, and by 30 minutes single nucleosomes are being further disassembled into sub-nucleosomes and free DNA, which continues through four hours.

Overall, this analysis indicates that the first stage of Enhancer Resetting involves a rapid (e.g. in minutes) decrease in nucleosome density at enhancer chromatin which corresponds to a change in both the kinetics of transcription factor binding and the ability of Tn5 to access the DNA. This initial change then gives way to a restoration of TF binding kinetics, while nucleosome occupancy continues to decrease.

### Chromatin remodeller activity controls re-establishment of transcription factor binding kinetics during Enhancer Resetting

The acute decrease in nucleosome density we see upon ERK activation reflects major changes in chromatin organisation at these key regulatory regions. We therefore asked whether chromatin remodellers play a role in coordinating the signal response. NuRD is a chromatin remodelling complex known to increase nucleosome density in a wide variety of systems^21,40,41^, and is required for ES cells to successfully undergo lineage commitment^17,20,42^. In contrast, while the BAF complex is also essential for lineage commitment^43,44^ it acts to reduce nucleosome density at regulatory regions^45–50^. We therefore hypothesised that these remodellers could be important in the enhancer resetting process.

To test the role of BAF in Enhancer Resetting we created a cell line in which the auxin inducible degron (AID) was fused to the C-terminus of the core, ATPase component BRG1 (encoded by the *Smarca4* gene) to create an inducible depletion system. Homozygous deletion of *Smarca4* leads to cell death in mouse ES cells^43,44^ however addition of auxin to the culture media of BRG1-AID cells resulted in depletion after 4 hours of treatment, allowing us to probe function of BRG1 in response to ERK signalling (Figure S4A). Using FA fixation and ChIP qPCR as an assay for Enhancer Resetting we subjected cells to BRG1 depletion and asked whether ChIP signal for NANOG was reduced after 8 minutes of ERK activation. Western blotting confirmed that there was no change in kinetics of the ERK signal response after BRG1 depletion (Figure S4B). Although BRG1 depletion results in reduced levels of NANOG at enhancers in 2iLIF, a further decrease in ChIP signal was seen after 8 minutes of ERK activation at a *Bmp4* enhancer with no sign of recovery after 4 hours (Figure 6A). This indicated that BRG1/BAF may not be required for the widespread mobilisation of transcription factors seen in the first stage of Enhancer Resetting. To more acutely regulate remodeller activity, we inhibited BRG1 using a small molecule inhibitor, BRM014^51^ for one hour prior to FA-fixation and NANOG ChIP (Figures 6B, S4B). Even where NANOG ChIP signal was low in the starting condition after BRG1 inhibition, a further decrease was seen 8 minutes after ERK activation (Figure 6B), confirming that BAF’s nucleosome remodelling activity is not required for the extensive changes in TF binding seen in Enhancer Resetting. While transcription factor enrichment is normally restored from 30 minutes of Enhancer Resetting, no such restoration is seen after BRG1-depletion (Figure 6A), consistent with the well demonstrated role of BRG1/BAF in facilitating transcription factor binding^45,46^.

**Figure 6.**
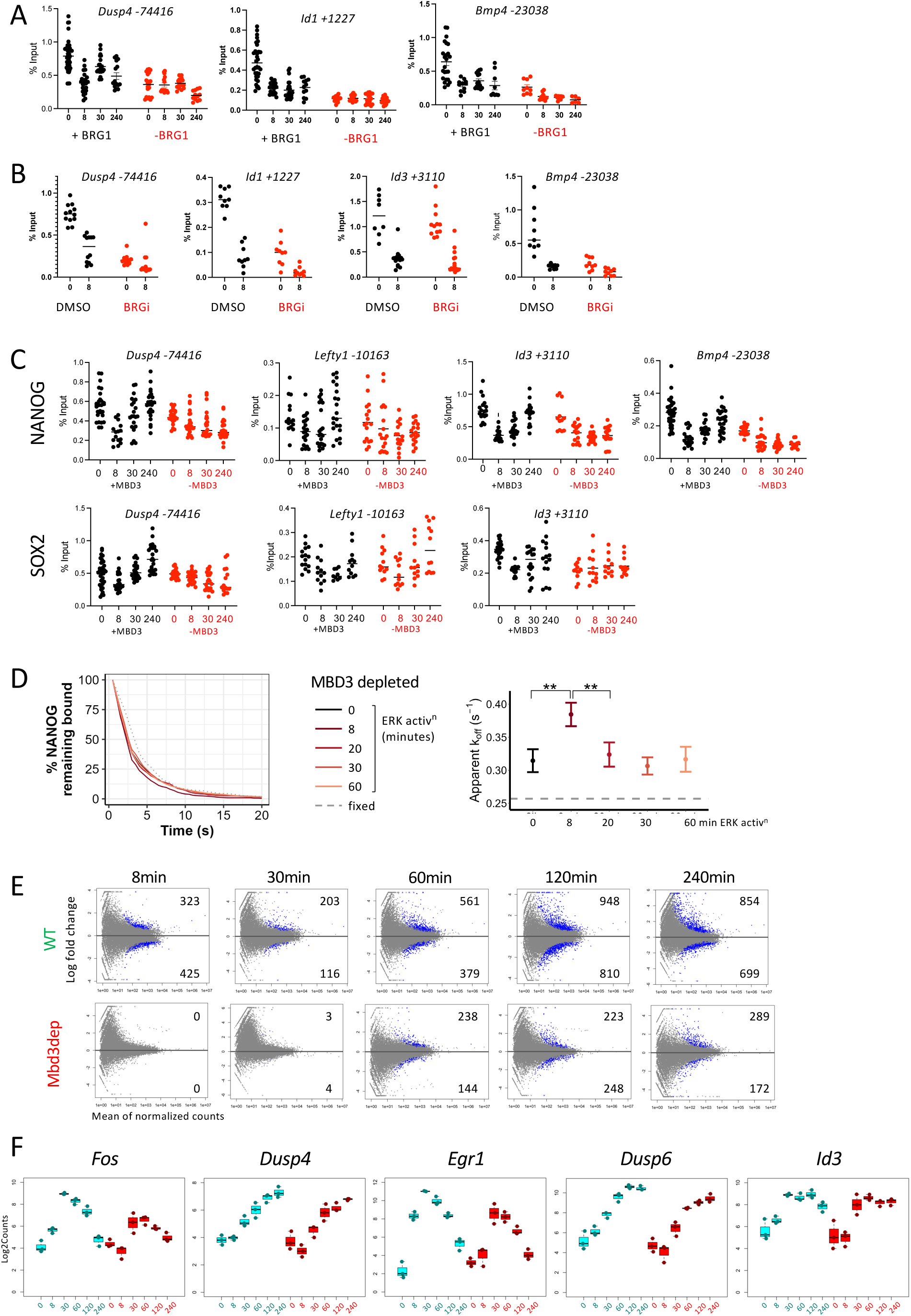
Chromatin remodeller activity is dispensable for the initial change in TF binding behaviour but is required for re-establishment and transcriptional response. **A**. NANOG ChIP-qPCR from FA-fixed samples. Data plotted shows the peak of NANOG enrichment at indicated sites before (black) or after 4 hours of BRG1-AID depletion (red). All replicates are plotted, with the line representing the mean. **B**. As in A but after either 1 hour pre-treatment with DMSO (black) or BRM014 (BRGi, red). **C.** As in A but either before (black) or after 60 minutes of MBD3-AID depletion prior to ERK activation time course (red). **D**. Florescence survival curves for NANOG-HALO localisations in MBD3 depleted cells (left) were used to calculate apparent dissociation rates (k_off_) (right) as in Fig. 3D. N = 4 biological replicates. **E**. Scatterplots indicating genes significantly changing expression (indicated as blue spots) relative to 2iL conditions in undepleted cells (top) or after MBD3 depletion (bottom) across the ERK activation time course. The numbers of significantly increasing or decreasing genes are indicated above or below the horizontal lines, respectively. **F**. RNA-seq expression profiles for indicated genes in undepleted (blue, left) or MBD3 depleted cells (red, right) across the ERK activation time course, time indicated in minutes on x-axes. Each data point is shown, and boxplots represent gene expression across three replicates. Boxes show the interquartile range (25th–75th percentile) with the median indicated by the centre line.

To enable us to test the function of NuRD in the ERK response we created two different NuRD inducible depletion ES cell lines. Firstly, both *Mbd3* alleles of an *Mbd2*-null ES cell line were targeted to express MBD3 with C-terminal fusion to AID^52,53^. MBD2 and MBD3 are mutually exclusive within NuRD but show partial redundancy^54–56^. These proteins link the remodelling and deacetylation subunits of the complex, such that loss of MBD3 in an *Mbd2*-null background results in complete dissociation of NuRD without affecting protein levels of the other subunits in the short term^53^ (Figure S4C). Secondly, ES cells were made in which both *Chd4* alleles were targeted to express a CHD4-FKBP fusion protein. This allows us to deplete CHD4 acutely upon addition of dTAG to the culture media^57^. CHD4 is the chromatin remodelling subunit of NuRD in ES cells but it can also function outside of the complex^58,59^, while MBD3 is only found within NuRD. Both *Mbd2^-/-^:Mbd3-AID* and *Chd4-FKBP* ES cell lines showed normal morphology and growth in the absence of Auxin or dTAG, respectively. Treatment of lines with Auxin or dTAG resulted in depletion of MBD3 or CHD4, respectively, from both the chromatin and nucleoplasm within 60 minutes^53^ (Figure S4C,D). In both cases, cells displayed appropriate ERK response (Figure S4E) and remained viable for ∼24 hours, after which they ceased proliferating and eventually died by apoptosis^53^.

ChIP-qPCR on FA-fixed, MBD3- or CHD4-depleted chromatin for NANOG or SOX2 showed reduced protein enrichment 8 minutes after ERK activation indicative of the first stage in Enhancer Resetting (Figures 6C, S4F). Furthermore, single molecule imaging showed an ERK-dependent increase in dissociation rate for NANOG in MBD3-depleted cells, comparable to that seen in wild type cells (Figure 6D). The ERK-induced increase in TF dissociation rate was therefore independent of NuRD activity. However, NuRD-depleted cells failed to show restoration of TF enrichment levels after this initial loss, indicating that NuRD is critical for the later stage of Enhancer Resetting.

We next asked whether NuRD’s chromatin remodelling activity contributes to the chromatin changes we observed upon ERK induction. NuRD activity is known to increase nucleosome density at enhancer sequences^21,40,41,49^ and this is reflected in a decrease in multinucleosome sized fragments (>300bp) and an increase in smaller sized fragments (<300bp) corresponding to monosomes, partial nucleosomes and free DNA at NANOG-, SOX2- and ETV5-bound sites after 1 hour of MBD3 depletion (Figure 5D). Depletion of MBD3 prevented both the dynamic changes in chromatin organisation and the changes in Tn5 integration frequencies induced by ERK activation in undepleted cells (Figures 5D, S3). This demonstrates that NuRD activity is required for the dynamic ERK-induced nucleosome remodelling characteristic of Enhancer Resetting.

NuRD activity is required for both the dramatic chromatin remodelling events and the re-establishment of transcription factor binding kinetics seen during enhancer resetting. To determine what consequences this has upon how the cell responds to ERK signalling, we next assessed how NuRD depletion impacts the transcriptional response to ERK activation. Nuclear RNA was collected from cells subjected to ERK activation after 1 hour of auxin-induced depletion of MBD3 and sequenced. While MBD3-depleted cells showed a clear transcriptional response to ERK activation, the magnitude of response was muted when compared to undepleted cells (Figures 6E; S4G). In the presence of NuRD activity ERK activation results in an immediate transcriptional response, with 325 and 425 differentially up- and down-regulated genes detected after 8 minutes respectively (Figure 6E). By contrast there were no significantly differentially expressed genes after 8 minutes in MBD3-depleted cells. While some genes show delayed activation in the absence of NuRD activity (e.g. Cluster 2, Figure S4G), those normally induced within 8 minutes (Cluster 5) are not induced at all in MBD3-depleted cells. This behaviour was confirmed using 4SU-labelling of nascent transcripts, with and without 1 hour pre-depletion of MBD3 (Figure S4H). Plotting the expression profiles of known ERK responsive genes clearly shows that there is both a delay and a reduction in amplitude of transcriptional response in the absence of NuRD. (Figure 6F). These data show that in the absence of NuRD activity, cells retain the ability to recognise the ERK signal but fail to mount an appropriate transcriptional response, the consequence of which, when multiplied across many genes, is ultimately a failure of lineage commitment^17,20,60^. We therefore conclude that the activity of chromatin remodellers are required for the resolution of TF dynamics during enhancer resetting and this activity facilitates a robust and appropriate transcriptional response to ERK signalling.

### Enhancer Resetting allows for signal specific transcription factor switching

We propose that when the ERK signal is received by a cell, RNAPII that is held in a paused state at enhancers is released and that subsequent transcription through regulatory sequences, either directly or through subsequent changes in chromatin structure, disrupts transcription factor interactions. We show that chromatin remodelling activity is required for these loci to return to a stable state. We hypothesised that this initial change in TF mobility would increase the possibility of new, signal specific factors engaging their target enhancers which are subsequently stabilised by remodelling of the chromatin. In this scenario however, at loci where there is no change in transcription factor required, or if the extracellular signal is not sustained, the original transcription factor binding repertoire would be re-established. To test this hypothesis, we looked at loci where TF binding is destabilised at 8 minutes and asked whether there was evidence for TF switching.

We identified 16937 sites bound by NANOG in 2iLIF conditions that showed decreased FA-ChIP signal at the 4 hour ERK activation timepoint. These are therefore loci that undergo destabilisation of NANOG binding upon ERK activation. While SOX2 enrichment at these sites is reduced at 8 minutes, it returns to near pre-stimulation levels after four hours (Figure 7A). Both NANOG and ETV5, however, show reduced enrichment at this later time point (Figure 7A) indicating a change in transcription factor occupancy at these enhancers after ERK activation. To investigate whether these sites become occupied by an ERK-responsive transcription factors, we endogenously epitope-tagged the long isoform of ETV5. Although ETV5 is present in ES cells in 2iLIF conditions, a longer ETV5 isoform transcribed from an upstream promoter is rapidly induced upon ERK activation^26^ (Figure S5). We performed ChIP-seq for the tagged long ETV5 isoform using cells grown either in ERK inhibition (2iLIF) or after 4 hours of ERK activation and plotted the signal across the sites where NANOG enrichment is reduced upon ERK activation (Figure 7B).

**Figure. 7.**
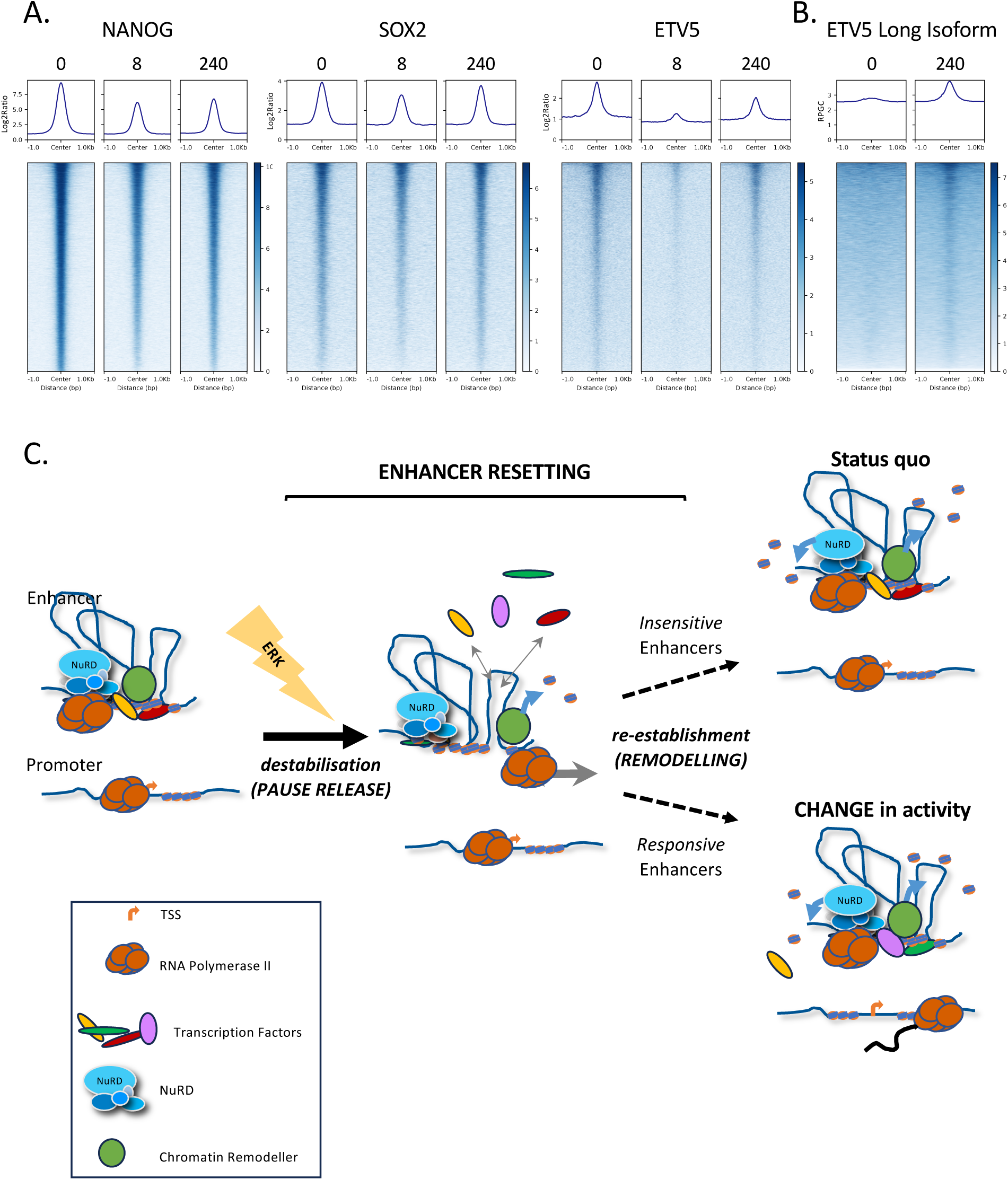
Enhancer Resetting enables efficient exchange of TFs in response to ERK signalling. **A**. ChIP-seq for NANOG, SOX2, and total ETV5 at indicated times after ERK activation is plotted across sites showing a significant loss of NANOG binding after 4h of ERK activation (N = 16937). **B.** ChIP-seq for ETV5 Long Isoform-3xFLAG as in A after 0 or 240 minutes of ERK activation. **C.** Schematic of Enhancer Resetting process in response to ERK signal activation. In steady state conditions before ERK signal is received (left side) transcription factors, remodellers and transcription machinery are all stably associated with regulatory regions. ERK signal results in the release of paused RNAPII which causes changes in chromatin architecture, destabilises TF interactions and thereby facilitates exchange of proteins at signal responsive regions (centre of figure). Establishment of a new repertoire of enhancer interacting factors is achieved through the action of the chromatin remodeller NuRD, returning the chromatin to a stable state and thereby resulting in the appropriate transcriptional response (right). At signal insensitive enhancers TF binding returns to original status with no change in transcription (“Status quo”) while ERK-responsive enhancers adopt an altered TF topology and a “CHANGE in activity”.

This showed that the long ERK-responsive ETV5 isoform was gained at regions losing both NANOG and total ETV5 binding during ERK activation. These data therefore support our hypothesis that Enhancer Resetting, induced by ERK activation, provides a window of opportunity for enhancers to switch their TF binding repertoire, and thereby rapidly initiate the signal-responsive transcriptional response (Figure 7C).

## Discussion

The receipt of an extracellular signal can rapidly change the gene expression programme and developmental trajectory of a cell. While signals can lead to long-term changes at the chromatin and gene expression levels, here we have focussed on determining how regulatory chromatin immediately responds to a change in the signalling environment to effect transcriptional change. By studying how the onset of an important signalling cascade activates enhancers in mouse ES cells at high temporal resolution, we have discovered a novel enhancer resetting event which precedes signal-induced changes in gene expression. We believe this has given us unprecedented insight into how transcriptional state changes occur. Enhancer Resetting is only detectable using methods that can identify this transient change in transcription factor kinetics. It will therefore have been missed in studies detecting TF enrichment using double-fixed ChIP, or those not including sufficiently short time courses. This novel event consists of two stages: an abrupt increase in the dissociation rate of transcription factors at enhancer chromatin (resulting in reduced residence times) within ten minutes of signal activation, and a recovery phase where transcription factor binding stability is restored over the next 1-2 hours and which correlates with the onset of transcriptional change. This event is coincident with an acute and transient change in chromatin accessibility globally at open chromatin regions. The first stage is dependent upon the ability of RNA Polymerase II to escape the paused state, while the recovery stage does not occur fully in cells depleted for either BAF or NuRD chromatin remodelling activity. We hypothesise that Enhancer Resetting creates a chromatin state that is permissive to exchange of transcription factors and thereby changes their probability of interacting with regulatory regions. We suggest that this mechanism of enhancer reconfiguration in response to intercellular signals is likely to be fundamental to developmental decisions.

How is the ERK signal first received by enhancer chromatin? That ERK pathway kinases can phosphorylate histone H3 at Serine 10 and Serine 28 is well documented^9,61,62^. These modifications have been described to facilitate the loss of repressive H3K9 and H3K27 methylation and the subsequent accumulation of histone modifications associated with active transcription, such as H3K27 acetylation^9,12,638^, consistent with the fact that we find H3K27 acetylation changes occur in the second stage coincident with transcription changes (Figure 2A, Figure S2A). DRB is known to mainly inhibit CDK9-mediated RNA Polymerase II phosphorylation, and its impact on the process reveals that release of transcriptional pausing is required in the initial stages. NELF activity (and therefore control of pausing) has been shown to be required for the transcriptional response to heat shock^64^, and NELFA, a key regulator of transcriptional pausing, is a direct target of ERK-mediated phosphorylation^65^. Phosphorylation of Serine 363 of NELFA not only occurs within 5 minutes of ERK activation^5^ but has also been shown to promote pause release^65^. While ERK mediated phosphorylation of NELF components could underlie the rapid release of paused RNA polymerase leading to a decrease in stable binding of TFs that we observe, ERK activation in ES cells results in rapid phosphorylation of a very large number of different proteins, several of which may be important for enhancer resetting to occur.

Enhancer Resetting does not convert enhancers from a silent to an active state. Rather, it occurs at regulatory regions with accessible chromatin that are already bound by transcription factors. Those enhancers ready to drive expression of ERK-responsive genes show a progressive increase in H3K27 acetylation after ERK activation, indicating that their activity has indeed been stimulated by onset of ERK signalling and subsequent Enhancer Resetting. Importantly, standard ATAC-seq analyses, where the abundance of ≤120bp reads is compared between sites, would not identify any changes in chromatin accessibility occurring during Enhancer Resetting. Mapping Tn5 integration frequency, however, allows for a more detailed view of accessibility levels in generally accessible chromatin. For example, as long as an accessible region has ≥2 Tn5 integrations within it, the region will generate an ATAC-seq signal. In contrast, measuring the number of integrations within a region gives a more quantitative indication of how accessible that chromatin is. Using this analysis, we find ERK activation causes an immediate but transient increase in accessibility across regulatory regions, which corresponds to the time when TFs show a change in binding kinetics.

We initially expected that the onset of Enhancer Resetting would be controlled by a chromatin remodeller but depletion or inhibition of several remodellers, including BRG1 and CHD4, all failed to prevent this initial step of enhancer resetting (Figures 6A,B, S4F). In contrast, preventing promoter escape of paused RNA polymerase II did prevent this early signal-induced change of TF binding kinetics (Figure 4). We propose that widespread release of Pol II from the paused state and subsequent transcription is sufficient to both destabilise TF binding and transiently increase Tn5 accessibility at regulatory elements. This is consistent with a previous study that identified changes in Mediator and RNA polymerase II enhancer association as driving factors in the transcriptional response to ERK^6^. In that case, however, TF binding was seen to change later in the response and was driven at the level of protein abundance, although TF behaviour was not assayed at very early timepoints. Indeed, other studies showed that NANOG protein levels decrease from 6 hours of ERK signal activation^27^ whereas we see no evidence for changes in protein levels in the early timepoints used in this study (Figure 2C). The initial destabilisation of TF binding that we describe appears to be largely indiscriminate, placing active enhancers into a transient, permissive state for regulatory change. If the activating signal persists, this transcriptionally driven destabilisation could facilitate the exchange of transcription factors while the abundance of the signal-responsive TFs remains relatively low. This exchange could therefore begin to take place within minutes of the signal being received, preceding changes in TFs at the protein level which typically take place over the course of hours.

After this first phase, chromatin accessibility decreases again to pre-stimulation levels, while nucleosomes continue to be disassembled and TF binding returns to pre-stimulation kinetics. During this period some enhancers re-stabilise with a different repertoire of associated TFs, and we propose that this enables them to change behaviour and drive ERK-responsive transcription (Figure 7C).

Chromatin remodeller activity is therefore important to create the chromatin environment on which Enhancer Resetting occurs - cells lacking NuRD activity display increased chromatin accessibility in 2iLIF compared to undepleted cells, and signal induction does not cause any further increase in accessibility. Although NuRD activity is not needed for the initial ERK-induced change in TF binding kinetics, NuRD-depleted cells show neither the remodelling of chromatin structure nor the re-establishment of TF binding patterns which occur in undepleted cells. Given that NuRD acts to increase the density of intact nucleosomes and to clear fragile nucleosomes^49,53^, we propose that NuRD activity, as well as that of BAF and probably other chromatin remodellers, is important to create the chromatin environment in which Enhancer Resetting can occur.

A widespread, sudden release of paused RNA polymerase, which we find is required for the initial destabilisation of TFs, could have a profound impact on enhancer chromatin. Pausing has been shown to be influenced by the presence of a very stable +1 nucleosome. In our current model, we suggest that ERK activation leads to release of pausing through phosphorylation and activation of pTEFb which would be sufficient to overcome this physical barrier to transcription. However, chromatin remodellers such as BAF and NuRD have opposing activities in controlling nucleosome occupancy at the TSS^41,66^ meaning that both their activities would be required to re-establish the nucleosomal architecture at regulatory regions once a new transcriptional program has been established. This could explain why both BAF- and NuRD-depleted cells are unable to fully carry out Enhancer Resetting in response to receipt of ERK signal, leading to a failure to mount a full and robust transcriptional response, culminating in developmental failure^20,43,44,60,67^.

Enhancer Resetting which we describe here enables rapid and efficient cellular response to ERK signalling in mouse ES cells allowing regulatory regions which are already bound by TFs to rapidly change their status. There is evidence that this mechanism may also occur in other cell types. A 2020 study from the Reyes group showed the induction of similarly widespread increases in chromatin accessibility at the majority of enhancers when Tgfß signalling was activated in mammary gland epithelial cells^68^. This effect occurred irrespective of the enhancers’ association with signal-responsive genes and exhibited similar kinetics to those seen in our current study. We suspect this could also reflect Enhancer Resetting in different cells, and in response to a different signalling system. Whether Enhancer Resetting is a general response to all signals or just some (or all) signals mediated by phosphorylation will be an interesting area for future investigation.

### Limitations of this study

Here we have studied the impact of ERK activation in mouse ES cells maintained in the presence of an ERK inhibitor. While going from ERK inhibition to ERK activation in naïve ES cells provides a simple, controllable system for careful molecular and biochemical dissection of the signalling response, most ERK-dependent developmental decisions will involve changes in levels, rather than a binary change in ERK activity^3^. We speculate that as negative feedback within the ERK pathway greatly dampens steady-state signalling^4^, receipt of additional input (e.g. a growth factor) stimulating the pathway could have a dramatic effect, involving Enhancer Resetting, in a developmental context. We propose that this is a general mechanism used by cells to rapidly re-configure enhancers to direct a signal-responsive transcriptional programme.

We identify and investigate the nature of Enhancer Resetting in naïve mouse ES cells. The very defined culture conditions used for mouse ES cells allowed us to methodically map the chromatin response to ERK activation and identify Enhancer Resetting. Now that we have identified this mechanism it will be possible to investigate how signal responses are mediated in other, more heterogeneous cell types with less defined culture conditions.

The precise role of BAF in the restoration phase of Enhancer Resetting is not clear. Inhibition or depletion of BRG1 results in a rapid reduction in binding of some transcription factors to their targets, and a loss of chromatin accessibility at enhancers^46,50,69^. The lack of a recovery phase in BRG1-depleted cells may therefore simply be due to a global dampening of NANOG binding in the absence of BAF activity. Any subsequent transcriptional changes could be due to Enhancer Resetting but could also be due to the global reduction in TF binding.

## Methods

### Key resources table

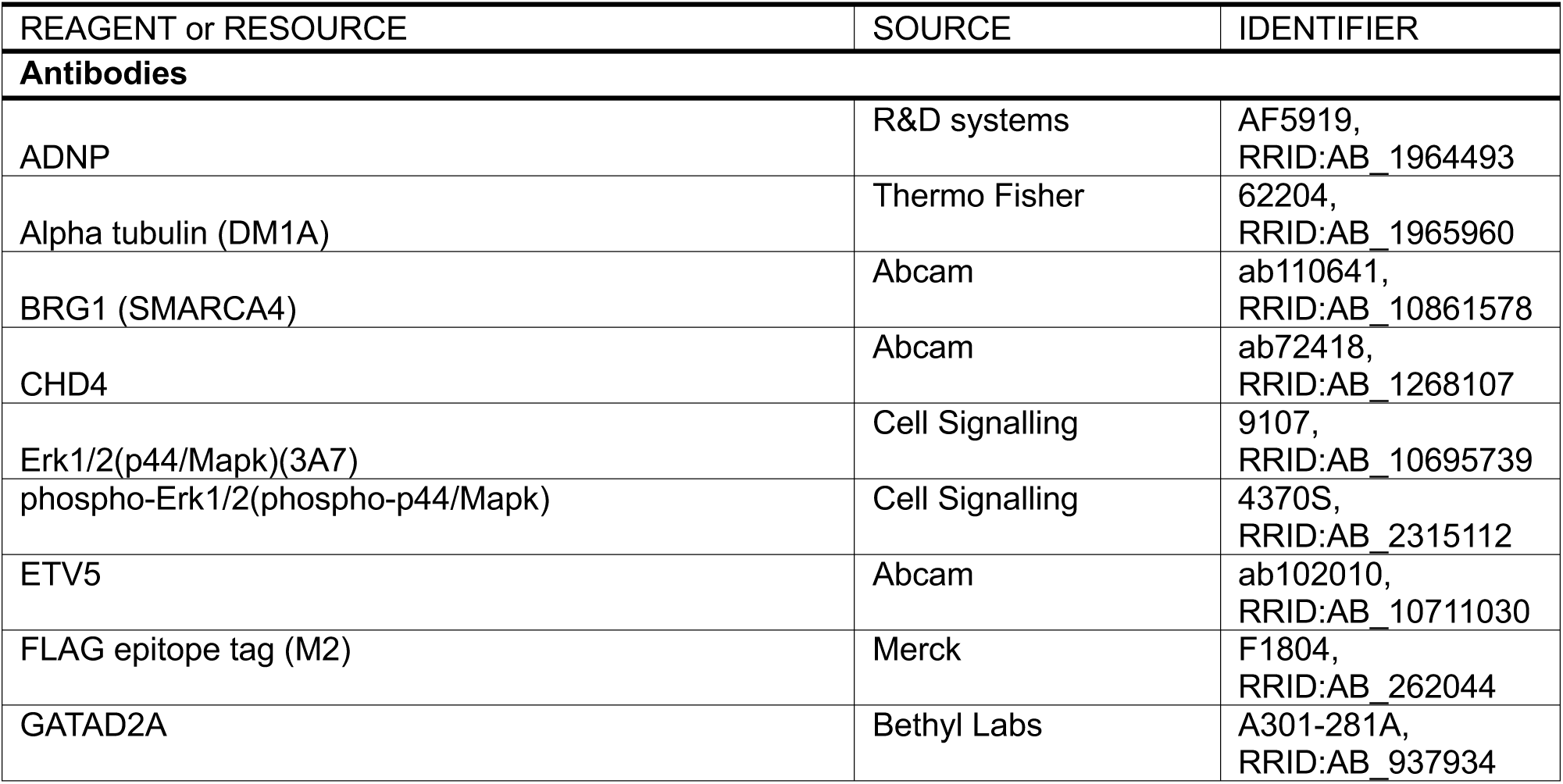
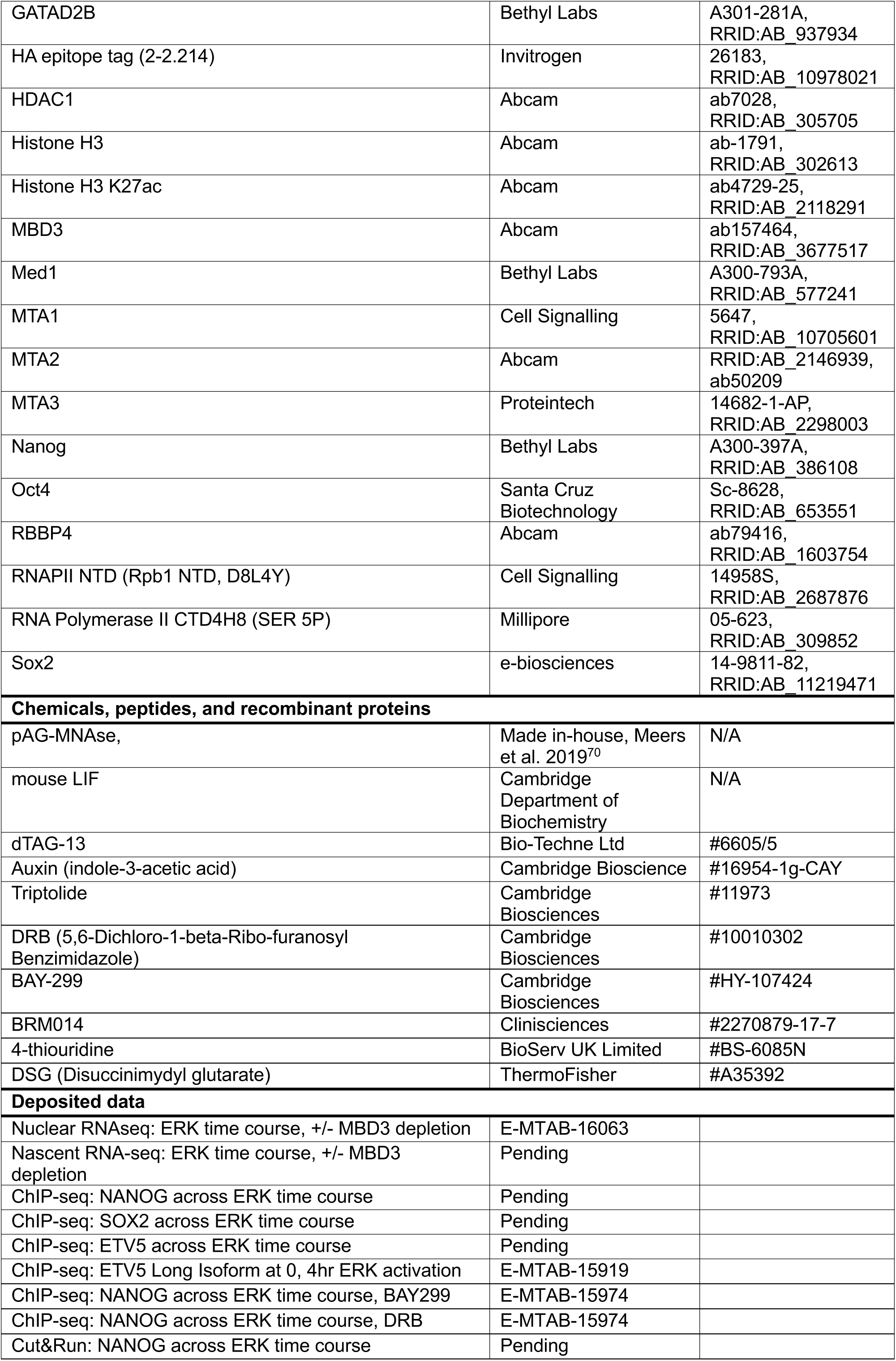
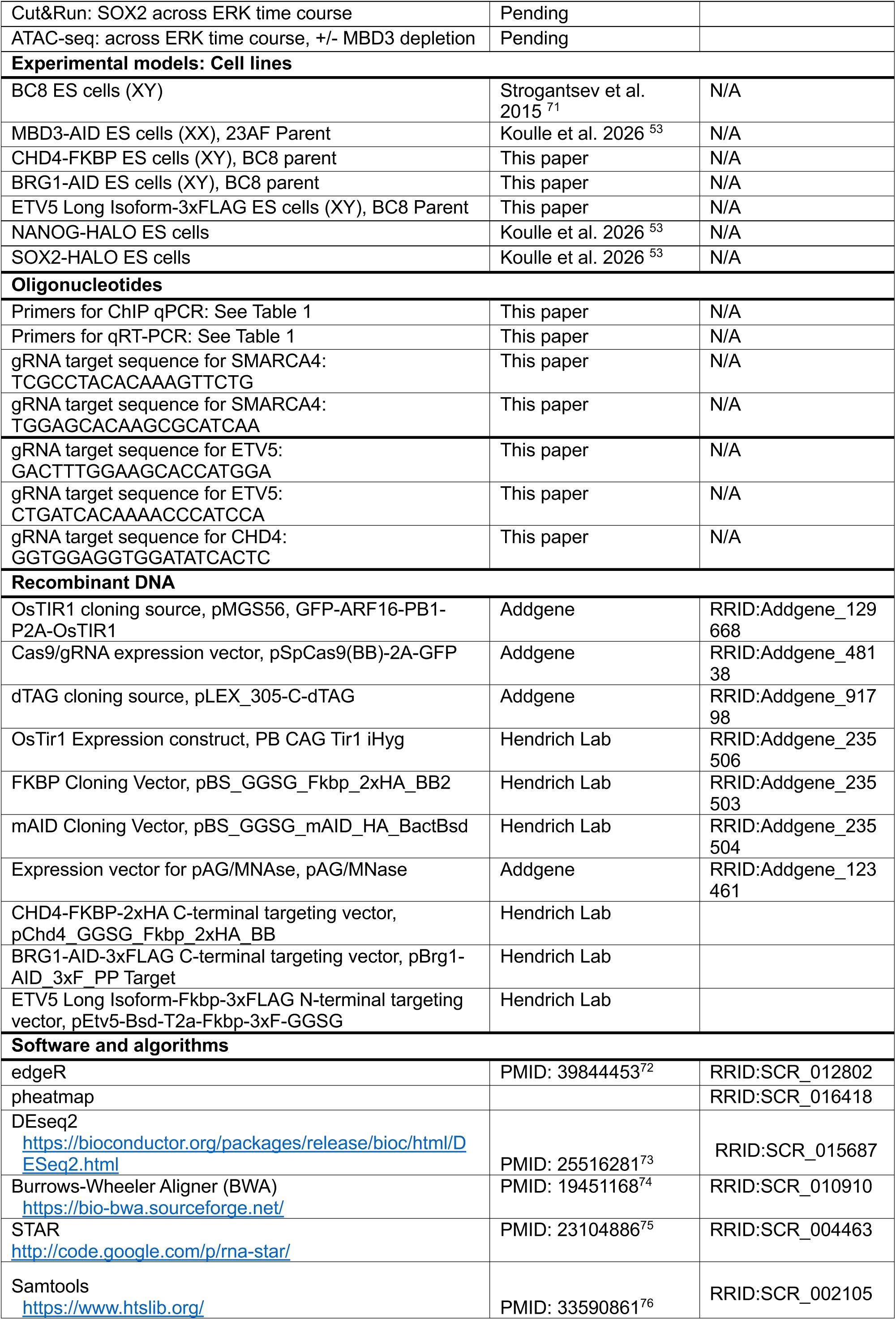
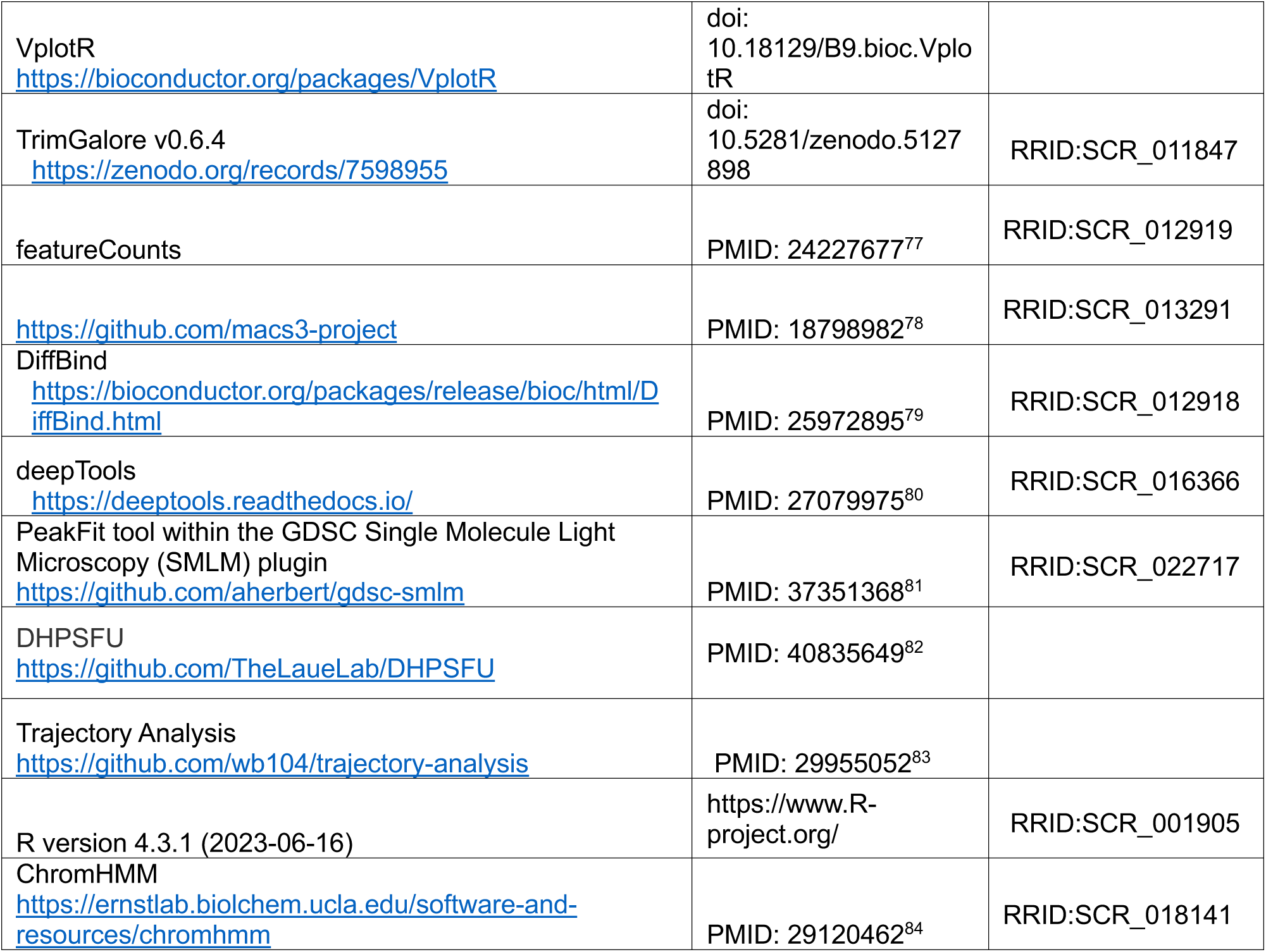

### Cell culture

Mouse ES cells were cultured on gelatinised plates in 2iL conditions (N2B27 supplemented with recombinant mouse LIF, 1 µM PD0325901 and 3 µM CHIR99021) as described^24^. Cell lines were regularly PCR genotyped and tested for mycoplasma contamination. ERK activation was carried out by replacing culture media with pre-equilibrated 1iL media (N2B27 supplemented with mouse LIF and 3 µM CHIR99021 only).

Depletion of degron tagged proteins was carried out by addition of specific small molecules with DMSO controls carried out in parallel. AID tagged proteins were depleted by addition of 0.5 mM auxin (indole-3-acetic acid) and FKBP tagged proteins were depleted using 0.5 µM dTAG-13 for the times indicated.

Chemical inhibition of Brg1 was carried out by addition of 10 µM BRM014 to culture media for 1 hour prior to start of time courses, and inhibitor maintained throughout experiments.

Inhibition of RNAPII transcription cycle was carried out by addition of the following inhibitors for 1 h prior to start of time courses, and inhibition maintained throughout: Triptolide at 1 µM; DRB (5,6-Dichloro-1-beta-Ribo-furanosyl Benzimidazole) at 100 µM; BAY-299 at 5 µM.

Cell lines used in this study are listed in the Key Resources Table. The BRG1-mAID-3xFLAG cell line was made in BC8, an F1 hybrid ES cell line from a C57Black/6 and Mus castaneus cross (40, XY); a gift from Anne Ferguson-Smith (Cambridge) ^71^. This cell line was first transfected with a PiggyBac construct to constitutively express OsTir1 (created using pMGS56); a gift from Michael Guertin (Addgene plasmid # 129668)) linked to Hygromycin resistance. Cells were subsequently cultured under Hygromycin selection to maintain OsTir1 expression. CHD4-FKBP and ETV5-3xFLAG cell lines were made in unmodified BC8 cells. Cells were lipofected with targeting constructs and an appropriate gRNA/Cas9 expression vector (made in pSpCas9(BB)-2A-GFP (PX458); a gift from Feng Zhang; Addgene plasmid # 48138). gRNA target sequences and targeting vectors are listed in the Key Resources Table. Correctly targeted clones were transiently transfected with Dre recombinase to remove ROXed drug selection cassettes, if present.

### Western Blots

Western blots for ERK activity were carried out on whole cell extracts by harvesting cells directly in sample buffer. Equal loading was confirmed by probing for total ERK and alpha-tubulin and ERK activation by phosphorylated ERK protein.

Levels of nuclear proteins were assessed by western blotting of either total nuclear extracts, or of isolated chromatin/nucleoplasm fractions^53,85^. Antibodies used are listed in Key Resources Table.

### Single Molecule Imaging

Long-exposure single molecule tracking (SMT) was carried out as described^53^, with the following adaptations. Trajectories were assigned a time-point in the ERK time-course based on when the first frame of the trajectory was detected in its respective SMT video. The dataset was then split into groups that represented the 8 min and 30 min ERK activation timepoints.

Trajectories were assigned to each of these two conditions using a window of 5-6 min centred around the exact timepoint.

Control samples were fixed in 4% Pierce™ formaldehyde (v/w) (Thermo Fisher, Cat. No. 28908) in PBS for 10 minutes at room temperature, rinsed twice with PBS and stored in PBS. Samples were imaged in identical pre-warmed culture media and under identical conditions to live-cell samples.

Short-exposure SMT was carried out with the same microscope set up as long-exposure imaging, with a frame rate of 50-100Hz. Short-exposure trajectories were segmented into confined and unconfined sub-trajectories using the 4P-algorithm^33^. Bound fractions were calculated for a sample as the number of trajectory frames assigned as being confined, divided by the total number of trajectory frames.

### RNAseq

Nuclear RNA was prepared according to Tellier et al 2022^86^ with minor adjustments. In brief, cells were grown to 70% confluency on 10 cm plates and subjected to ERK activation as detailed. Cells were washed twice with ice-cold PBS and harvested by scraping in ice-cold PBS. After centrifugation at 500 xg for 5 minutes at 4°C, cell pellets were resuspended in 1 ml Lysis Buffer (10 mM Tris-HCl pH 8.0, 140 mM NaCl, 1.5 mM MgCl_2_, 0.5% NP40) and pellets collected after further centrifugation at 1000 xg for 3 minutes at 4°C. The resulting pellet was resuspended in 1 ml Lysis Buffer, transferred to a round bottomed tube and 100 µl Detergent Solution (3.3% sodium deoxycholate, 6.6% Tween-20) added dropwise while vortexing slowly. Samples were transferred to 1.5 ml tubes and centrifuged at 100 xg, 5 minutes at 4°C. Nuclear pellets were then resuspended in 300 µl Trizol and RNA extracted according to manufacturer’s protocol.

Nuclear RNA sequencing reads were aligned to the mouse reference genome (mm10/GRCm38) using STAR^75^ with parameters suitable for spliced read alignment. Gene-level read counts were generated from aligned reads to produce a count matrix for downstream analysis. Prior to differential expression analysis, genes with low expression across samples were filtered out. Differential gene expression analysis was performed using the DESeq2 package in R^73^. Read counts were normalized using DESeq2, and statistical testing was performed by comparing each time point to the 2i/LIF baseline condition. Adjusted p-values were calculated using the Benjamini–Hochberg method^87^

For visualization and clustering analyses, normalised counts were transformed using the regularised log (rlog) transformation implemented in DESeq2. Heatmaps of gene expression across samples were generated using the pheatmap package in R with hierarchical clustering applied to both genes and samples. To identify groups of genes exhibiting similar temporal expression dynamics during ERK activation, K-means clustering was performed on rlog-transformed expression values across all time points. To visualize cluster-specific transcriptional responses, expression changes relative to the baseline condition were calculated as Δlog₂ expression (log₂ fold change relative to 2i/LIF). The mean Δlog₂ expression of genes within each cluster was calculated and plotted across the ERK activation time course.

Sequencing of nascent RNA was carried out by metabolic labelling of nascent transcripts by addition of 500 µM 4-thiouridine to culture media for 5 minutes, starting and ending 2.5 minutes either side of each ERK activation timepoint. RNA was processed and purified according to Radle et al 2013^88^.

Raw RNA-seq reads were aligned to the mouse reference genome (mm10/GRCm38) using the Burrows-Wheeler Aligner (BWA) ^74^. To quantify nascent transcription, reads mapping to the first 1 kb downstream of annotated transcription start sites (TSS) were counted using featureCounts^77^. Differential transcription analysis was performed using the edgeR package^72^ in R. Count data were normalized within the edgeR framework, and statistical testing was used to identify genes showing significant changes in nascent transcription relative to the 2i/LIF baseline condition. Normalized expression values were visualized using heatmaps generated with the pheatmap package in R. Genes were grouped according to their transcriptional patterns using hierarchical clustering.

### ATACseq

Cells were grown on 15 cm plates, treated with auxin to deplete MBD3 protein where noted, before ERK activation time course. Due to the short timepoints, nuclei were harvested and frozen in the presence of DMSO at −70 according to Kaya-Okur et al 2020^89^. ATAC was carried out using Illumina reagents (#15027865) according to the manufacturer’s protocol. Sequencing was carried out by the CRUK Cancer Institute sequencing facility.

Sequencing reads were aligned to the mouse reference genome (mm10/GRCm38) using BWA. Alignment files were processed using SAMtools^92^ including conversion to BAM format, sorting, removal of duplicate reads, and removal of reads mapping to the mitochondrial genome. Chromatin accessibility patterns were analysed using the VplotR package in R. V-plots were generated to visualize fragment length distributions and read density across genomic regions of interest. Heatmaps of read density were produced, and visualization parameters were adjusted to mitigate saturation effects, enabling visualization of both high- and low-density fragment signals.

Normalized Tn5 integration site counts were calculated across regions of interest to quantify accessibility. Using the matrices generated for V-plot visualization, fragment density profiles were calculated for distinct fragment length classes corresponding to chromatin states: <80 bp (nucleosome-free fragments), 80–149 bp (sub-nucleosomal fragments), 150–300 bp (mono-nucleosomal fragments), and >300 bp (di-nucleosomal fragments). Normalized read densities for each fragment class were plotted across regions of interest to assess changes in chromatin accessibility and nucleosome organization.

### Cut and Run and Chromatin Immunoprecipitation

Cut&Run was carried out as described^70^. 100,000 live ES cells were used per reaction, with 10,000 drosophila SL2 nuclei added as a spike-in control. pAG-MNAse was made and purified in-house as described^70^. Plasmid pAG/MNase was a gift from Steven Henikoff (Addgene plasmid #123461). Library preparation was carried out in the CSCI Genomics facility.

For chromatin immunoprecipitation, cells were fixed on culture plates in 1% formaldehyde for 10 minutes at room temperature and fixation quenched by addition of glycine to 125mM. For double fixation, cells were first harvested in ice cold PBS before resuspension in 2mM DSG (Disuccinimydyl glutarate) in PBS and incubation for 50min at room temperature before fixation in 1% formaldehyde and quenching with glycine as above. Chromatin immunoprecipitation was carried out as described^21^ using 150 µg chromatin per IP and antibodies as listed in Table X. 15µg of Rat ES cell chromatin was included in each sample as spike-in. Rat ES cells were a gift from Austin Smith, Exeter^93^. Library preparation was carried out in the CSCI Genomics facility and sequenced at the CRUK Cancer Institute sequencing facility.

ChIP carried out in the presence of BAY-299 or DRB and that performed for the ETV5 Long Isoform included 15 µg Drosophila SL2 chromatin as spike-in, and library construction and sequencing were carried out by Novagene.

Sequencing reads from ChIP–seq and CUT&RUN experiments were processed using the same computational workflow. Raw reads were trimmed using TrimGalore v0.6.4 to remove adapter sequences and low-quality bases. Trimmed reads were aligned to the mouse reference genome (mm10/GRCm38) using BWA. Alignment files were processed using SAMtools, including conversion to BAM format, sorting, indexing, and filtering of low-quality reads.

Peak calling was performed using MACS2^78^ with corresponding input controls to identify transcription factor binding sites. Peaks were called independently for each sample using parameters appropriate for narrow peak detection. Differential binding analysis was performed using the DiffBind package^79^ in R. Consensus peak sets were generated for each transcription factor across all samples, and read counts were quantified across these regions. Counts were normalized using the DESeq2 framework implemented within DiffBind. Differential binding was assessed by comparing the 2i/LIF baseline condition to subsequent time points following ERK activation.

For visualization of genome-wide binding patterns, normalized coverage tracks were generated using deepTools^80^. BigWig files were produced using bamCoverage, and signal enrichment across genomic regions was visualized using computeMatrix, plotHeatmap, and plotProfile in R.

**Table 1.**
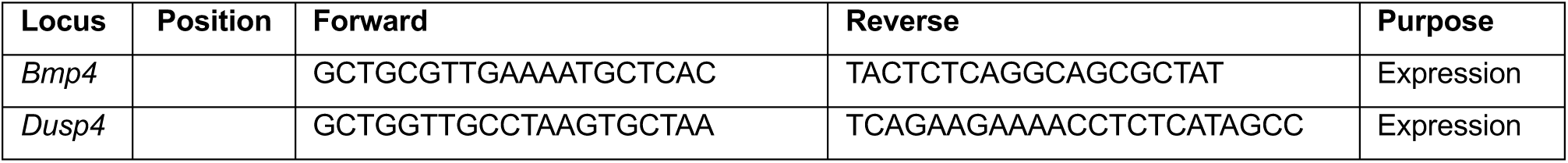
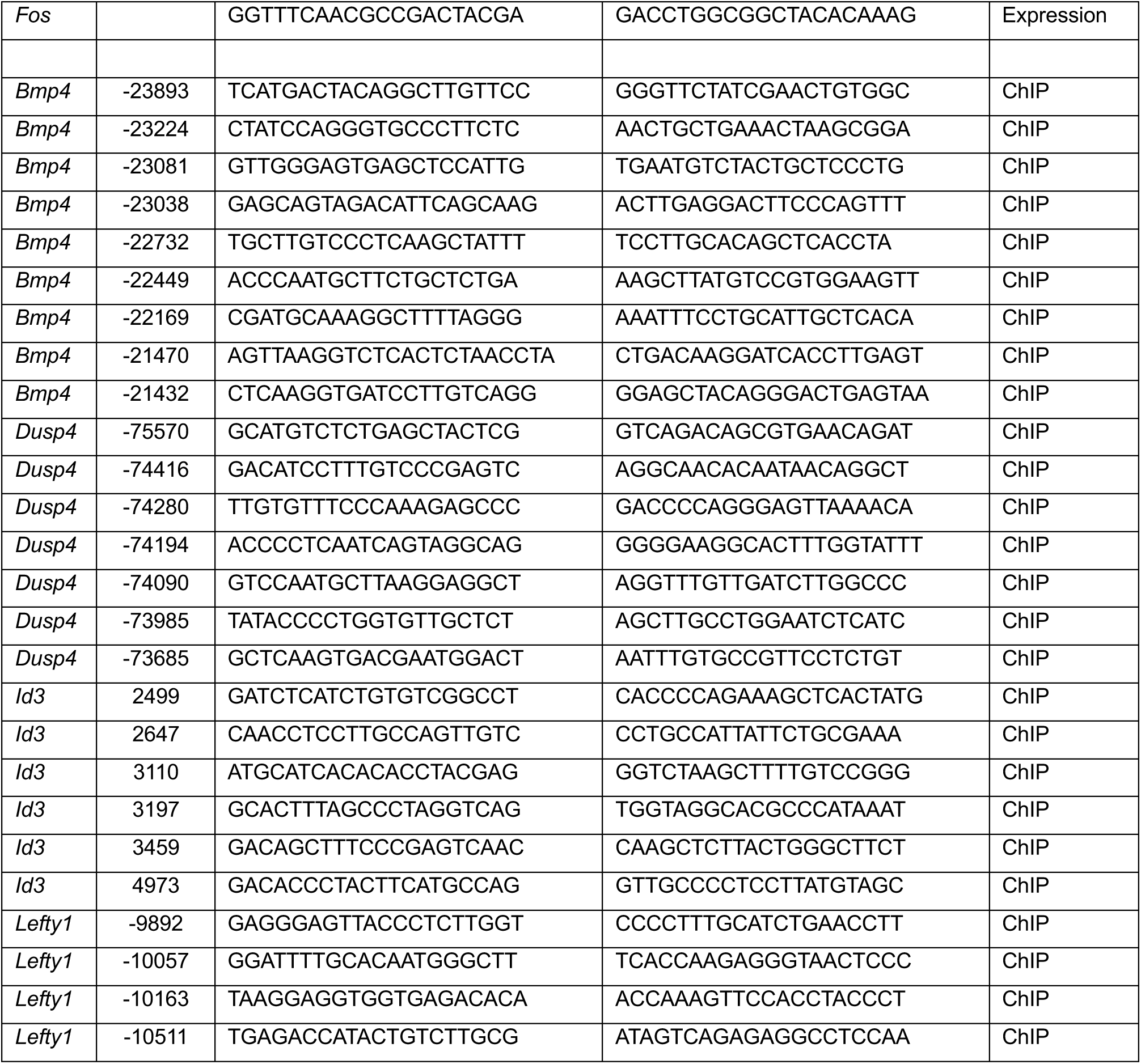
PCR primers used in this study. Position for ChIP-qPCR denotes distance of mid-point of PCR product from TSS.

## Acknowledgments

We are grateful to past and present members of the Hendrich and Laue Groups and to the Fragile Nucleosome online community for useful discussions. We are grateful to the CSCI Cell Culture and NGS facilities for expert support. We also thank the project and rotation students whose work has contributed to the idea of Enhancer Resetting over the years.

## Funding

This work was supported by grants from the Medical Research Council to B.D.H. (MR/R009759/1, MR/X018342/1 and MR/Y000595/1) and to E.D.L. (MR/P019471/1 and MR/M010082/1), from the Wellcome Trust to E.D.L. (206291/Z/17/Z), from the Isaac Newton Trust to B.D.H. (17.24(aa)), a PhD studentship from the Medical Research Council to D.S., and core funding from the Wellcome/MRC (203151/Z/16/Z) to the Cambridge Stem Cell Institute.

## Author Contributions

NR, AK, JB, RF, PG and BH created and validated ES cell lines; NR, BH, AK, OO, NM and RF generated and analysed qPCR data, NR, JB and BH generated sequencing data; RR, ML, and NR analysed sequencing data; DS and EDL conducted single molecule imaging experiments and analyses; RR developed bioinformatic tools and supervised analyses; NR, EDL and BH supervised the project and EDL and BH obtained funding. The manuscript was written by NR and BH with input from all other authors.

## Competing Interests Statement

The authors declare no competing interests.

## Materials Availability Statement

High throughput sequencing data generated in this project are available from Array Express with accession numbers indicated in the Key Resources Table. Plasmids created as part of this study are available from Addgene: https://www.addgene.org/Brian_Hendrich/. Cell lines can be requested from BDH.

## Notes

### Competing Interest Statement

The authors have declared no competing interest.

