## Supplemental Figures for "Chromatin remodelling enables enhancer resetting to facilitate the ERK transcriptional response"

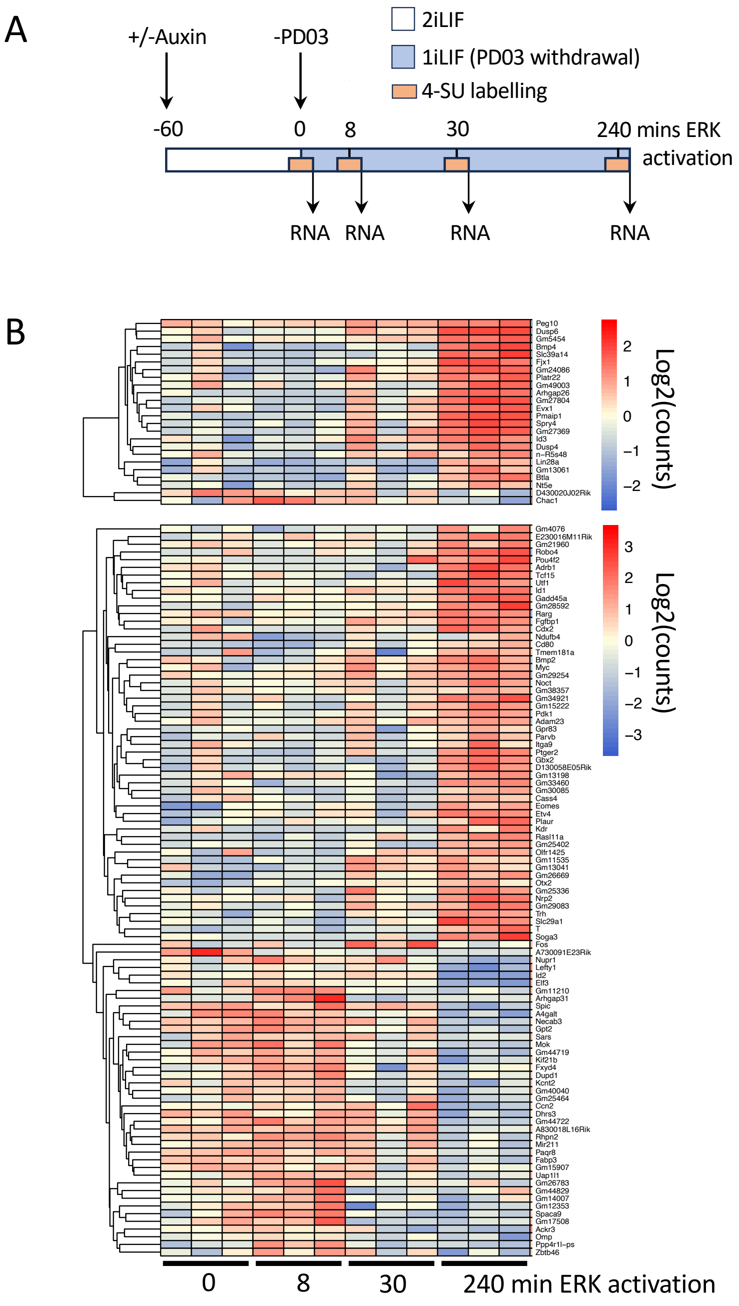


**Supplemental Figure 1. A.** Schematic of nascent RNA sequencing experiment. Cells were labelled with 4SU for five minutes spanning the indicated time points (orange boxes) prior to harvesting and RNA purification. ERK activation by withdrawal of the inhibitor PD0325901 (PD03) is indicated in blue. **B**. Heat map constructed for genes showing significant changes in nascent RNA sequencing across the ERK activation time course at indicated times (FDR ≤ 0.05). Data for each of three biological replicates is shown. Gene names are indicated at right.

Supp Fig. 2

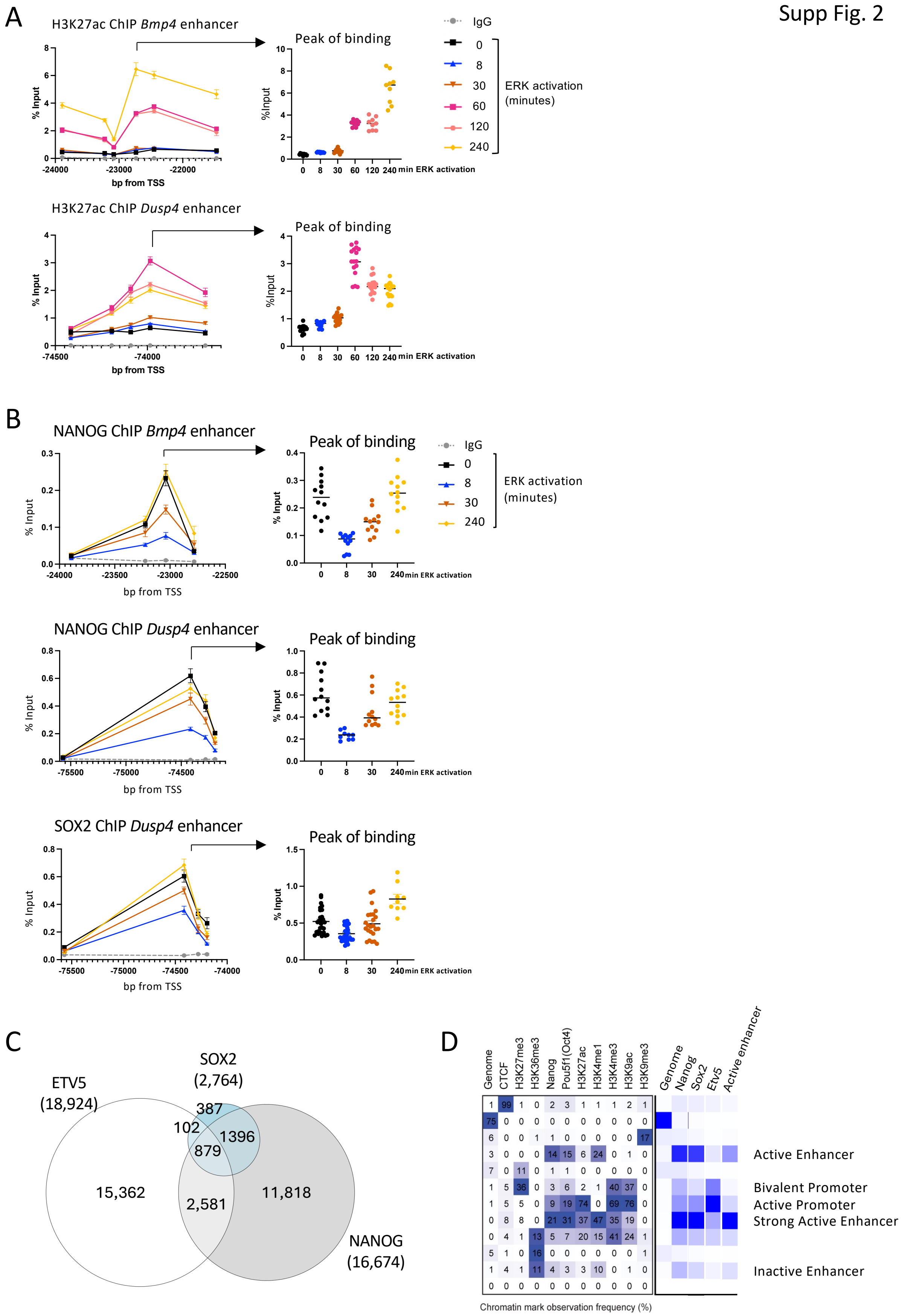


**Supplemental Figure 2**. **A**. ChIP-qPCR for H3K27Ac was performed at indicated times of ERK activation. Data are plotted across enhancers located near *Bmp4* or *Dusp4*, with x- axis labels indicating base pairs relative to the annotated TSS of each gene. For each plot all replicates from the peak of binding are re-plotted at right at indicated times of ERK activation. Line graphs show mean and standard error (N ≥ 3) while peak of binding plots show all replicate values and the horizontal line indicates the mean. **B**. As in A, but for NANOG or SOX2 across indicated enhancers. N ≥ 3. **C**. Venn diagram of peaks showing differential signal between 0 and 8 minute timepoints, identified in ChIP-seq data for indicated transcription factors. **D**. ChromHMM [84] results for ChIP-seq peaks for NANOG, SOX2, ETV5, or a set of active enhancers [21] .

Supp Fig. 3

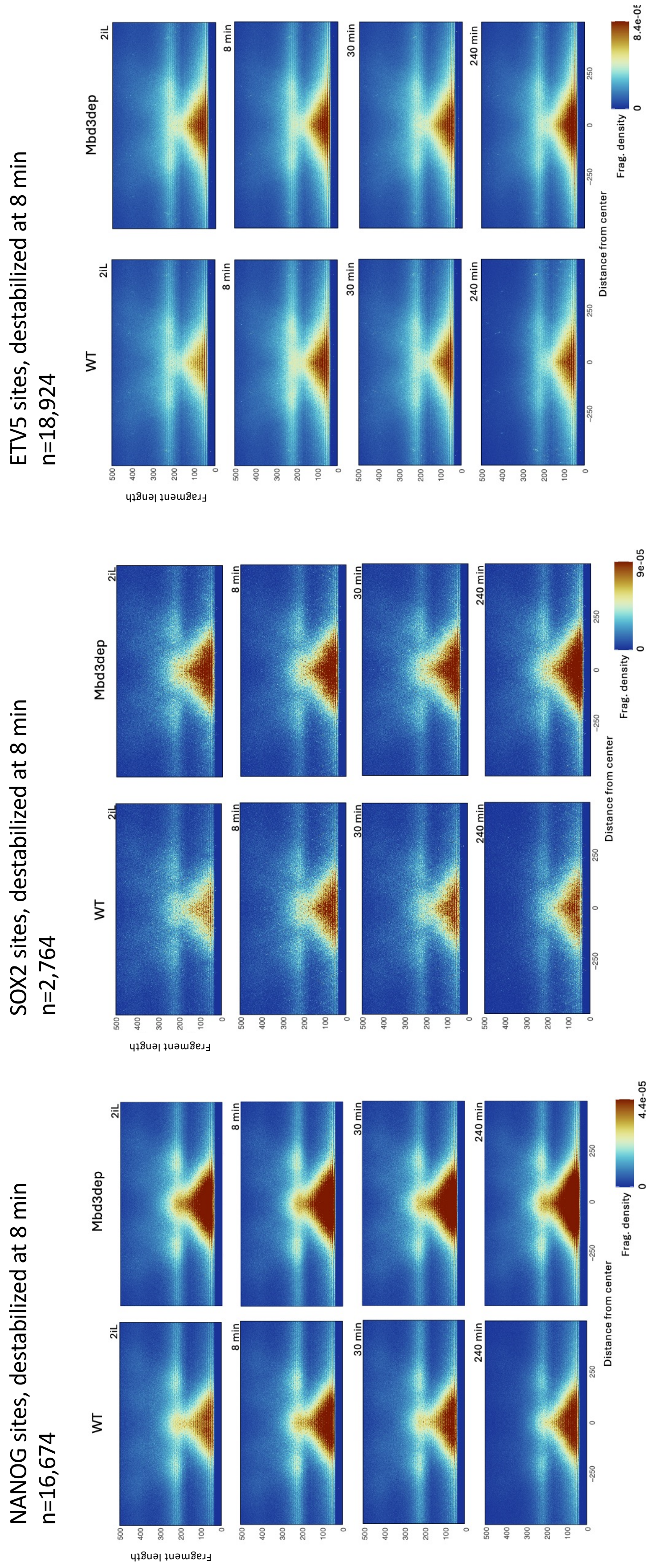


**Supplemental Figure 3.** Vplots as in Figure 5B for sites showing significant changes in enrichment for NANOG, SOX2 or ETV5 after ERK activation in wild type cells (left panels) or after MBD3 depletion (right panels). The data from Figure 5B are re-plotted here (left hand panels) for ease of comparison.

Supp Fig. 4

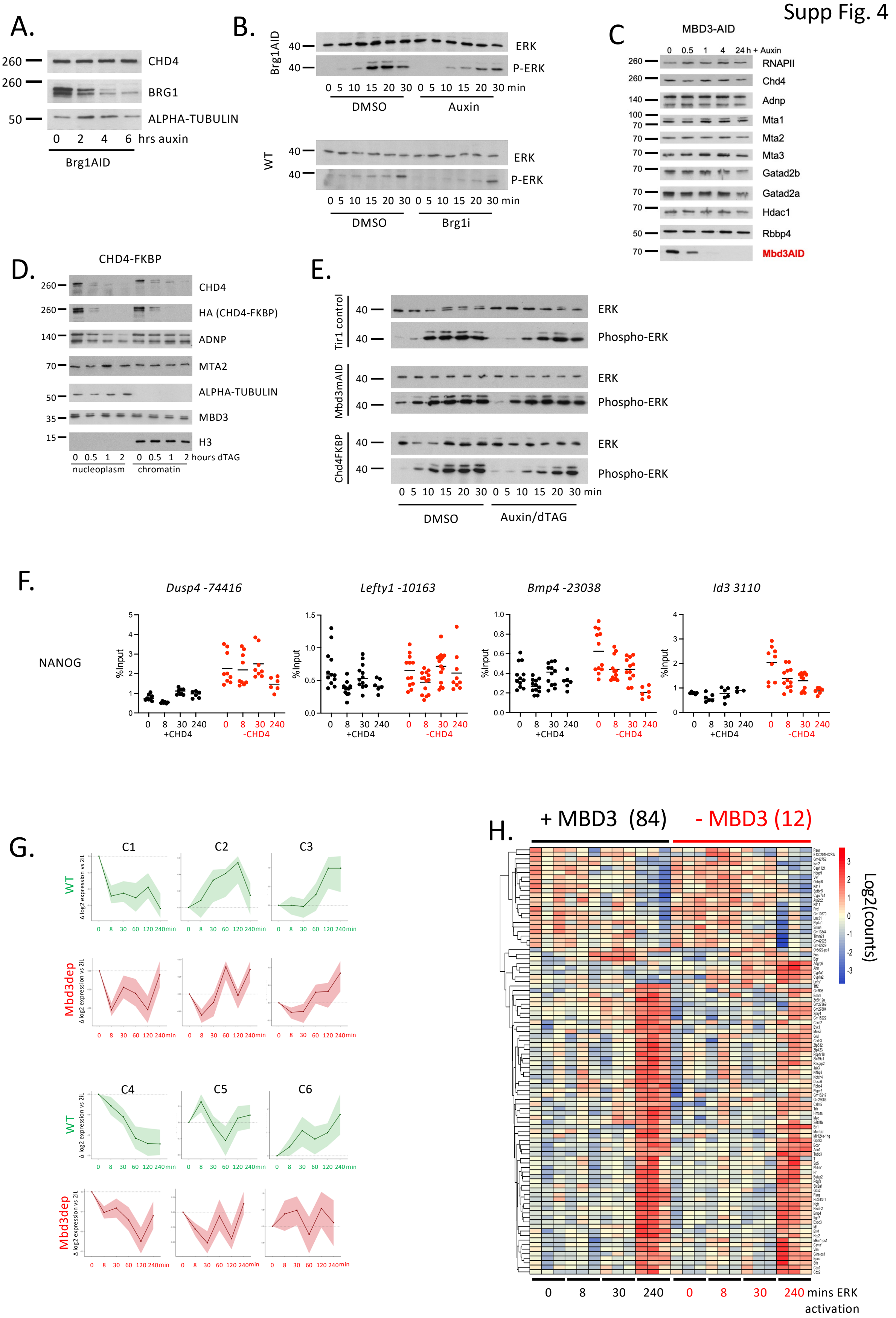


**Supplemental Figure 4. A.** Western blots showing auxin mediated depletion of BRG1 in BRG1AID line. Nuclear extracts from cells exposed to auxin for various times (noted on bottom) were probed for CHD4 (top), alpha-tubulin (bottom) which act as loading controls, and for BRG1.

Position of size markers in kDa is indicated on the left. **B.** Western blots to show kinetics of ERK response in BRG1 depleted cells (top) and in BRG1 inhibition (bottom). ERK activation was induced by removed of PD03 from culture media for times shown at bottom. Whole cell extracts from cells in the presence of DMSO control or either auxin or BRG1 inhibitor BRM014 as indicated were probed for total levels of ERK1/2 (top) as loading control, and for phospho-ERK (bottom blot of each set). Size markers are indicated in kDa. **C.** Auxin mediated depletion of MBD3AID. Western blots of nuclear extracts from cells grown in the presence of auxin for the times indicated probed for NuRD component proteins as indicated. RNAPolII acts as a loading control. Sizes are shown in kDa. **D.** Depletion of CHD4FKBP protein in response to dTAG. Nucleoplasm and chromatin fractions from cells exposed to dTAG for the times shown were probed for NuRD and ChAHP components as indicated, with alpha-tubulin and Histone H3 acting as loading controls for nucleoplasm and chromatin fractions respectively. Sizes are shown in kDa. **E.** ERK response in MBD3 and CHD4 depleted or control (Tir1) cells. As in B but cells were grown in the presence of auxin (MBD3AID and TIR1) or dTAG (for CHD4FKBP). **F.** NANOG ChIP-qPCR data at the peak of NANOG enrichment at indicated sites before (black) or 1 hours after CHD4-FKBP depletion (red). All replicates are plotted, with the line representing the mean. **G.** Cumulative behaviour of ERK responsive gene clusters identified in Figure 1C, in the presence (green) or absence of MBD3 (red). Line shows mean gene expression; shaded ribbon indicates the interquartile range (25th–75th percentile). Wild type profiles from Figure 1C are reproduced for ease of comparison. **H.** Heat map constructed for genes showing significant changes in nascent RNA sequencing across the ERK activation time course at indicated times (FDR 0.05), with wild type data reproduced from Figure S1B on left and MBD3 depleted cells on right. Data for each of three biological replicates is shown. Gene names are indicated at right.

Supp Fig. 5

A

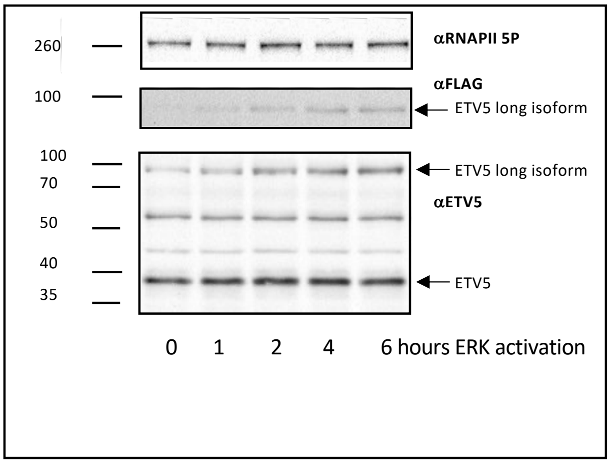


**Supplemental Figure 5.** Western blots of nuclear extracts exposed to ERK activation time course, time indicated at bottom. Blots probed for FLAG to detect induction of ERK specific ETV5 long isoform, and with antibody to ETV5 to below. Serine 5 Phospho-RNAPII acts as a loading control. Sizes are shown in kDa.
